# Identification of DLK1, a Notch ligand, as an immunotherapeutic target and regulator of tumor cell plasticity and chemoresistance in adrenocortical carcinoma

**DOI:** 10.1101/2024.10.09.617077

**Authors:** Nai-Yun Sun, Suresh Kumar, Yoo Sun Kim, Diana Varghese, Arnulfo Mendoza, Rosa Nguyen, Reona Okada, Karlyne Reilly, Brigitte Widemann, Yves Pommier, Fathi Elloumi, Anjali Dhall, Mayank Patel, Etan Aber, Cristie Contreras-Burrola, Rosie Kaplan, Dan Martinez, Jennifer Pogoriler, Amber K. Hamilton, Sharon J. Diskin, John M. Maris, Robert W. Robey, Michael M. Gottesman, Jaydira Del Rivero, Nitin Roper

## Abstract

Immunotherapeutic targeting of cell surface proteins is an increasingly effective cancer therapy. However, given the limited number of current targets, the identification of new surface proteins, particularly those with biological importance, is critical. Here, we uncover delta-like non-canonical Notch ligand 1 (DLK1) as a cell surface protein with limited normal tissue expression and high expression in multiple refractory adult metastatic cancers including small cell lung cancer (SCLC) and adrenocortical carcinoma (ACC), a rare cancer with few effective therapies. In ACC, ADCT-701, a DLK1 targeting antibody-drug conjugate (ADC), shows potent *in vitro* activity among established cell lines and a new cohort of patient-derived organoids as well as robust *in vivo* anti-tumor responses in cell line-derived and patient-derived xenografts. However, ADCT-701 efficacy is overall limited in ACC due to high expression and activity of the drug efflux protein ABCB1 (MDR1, P-glycoprotein). In contrast, ADCT-701 is extremely potent and induces complete responses in DLK1^+^ ACC and SCLC *in vivo* models with low or no ABCB1 expression. Genetic deletion of DLK1 in ACC dramatically downregulates ABCB1 and increases ADC payload and chemotherapy sensitivity through NOTCH1-mediated adrenocortical de-differentiation. Single cell RNA-seq of ACC metastatic tumors reveals significantly decreased adrenocortical differentiation in DLK low or negative cells compared to DLK1 positive cells. This works identifies DLK1 as a novel immunotherapeutic target that regulates tumor cell plasticity and chemoresistance in ACC. Our data support targeting DLK1 with an ADC in ACC and neuroendocrine neoplasms in an active first-in-human phase I clinical trial (NCT06041516).

## Introduction

Targeting cell-surface antigens with antibody-drug conjugates (ADCs) is a promising immunotherapeutic approach in oncology with recent FDA approvals across a diverse set of malignancies. Nonetheless, identification of new tumor-specific targets is imperative, especially for refractory adult metastatic tumors with few treatment options, and for less common malignancies, such as neuroendocrine (NE) neoplasms, with unique biological features.

One defining feature of neuroendocrine neoplasms, as the categorization of these cancers implies, is NE differentiation, characterized by high expression of a coordinated set of genes, including synaptophysin and chromogranin, routinely used as clinical diagnostic markers for these tumors. The Notch pathway is a major negative regulator of neuroendocrine differentiation and suppression of this pathway is common across neuroendocrine tumors^1^. While mechanisms of Notch pathway suppression in neuroendocrine cancers are not entirely clear, it is known that the cell surface Notch ligands such as delta-like 3 (DLL3) inhibit Notch pathway activation in normal development^2^. Moreover, as DLL3 expression is restricted to the brain but aberrantly expressed in many neuroendocrine cancers, DLL3 was an early immunotherapeutic target in small cell lung cancer (SCLC)^3^ and neuroendocrine prostate cancer^4^. While initial efforts to target DLL3 with an ADC failed, more recent efforts targeting DLL3 via T-cell engager strategies in SCLC have demonstrated remarkable success^5,6^ with recent approval by the FDA for treatment of relapsed SCLC.

To our knowledge, there has been no systematic effort to assess whether Notch ligands beyond DLL3 may or may not be targetable cell surface proteins in cancer. Therefore, in this work, we screened normal tissue and metastatic cancer datasets for expression of Notch ligands (DLL1, DLL3, DLK1, JAG1, JAG2) and uncovered DLK1 (delta-like non-canonical Notch ligand 1) as a candidate cell surface immunotherapy target protein. Moreover, we show that DLK1 is targetable by an ADC, particularly in the rare cancer adrenocortical carcinoma (ACC) in which DLK1 is highly expressed. Importantly, we find that DLK1 is a key driver of chemoresistance in ACC through maintenance of adrenocortical differentiation and expression of the drug efflux protein ABCB1 (MDR1, P-glycoprotein) thereby demonstrating an important biological function for this new immunotherapeutic target.

## Results

### DLK1 has limited normal tissue expression and high expression in multiple metastatic cancers including adrenocortical carcinoma

To assess whether Notch ligands could be suitable cell surface immunotherapeutic targets, we compared normal tissue expression of *DLL1*, *DLL4*, *DLK1*, *JAG1*, and *JAG2* with *DLL3* using the adult Genotype-Tissue Expression (GTEx) Portal^7^. As expected, expression of *DLL3* was restricted to the brain (Supplementary Fig. 1A). However, other Notch ligands (*DLL1*, *DLL4*, *JAG1*, and *JAG2*) were expressed across a wide span of normal tissues (Supplementary Fig. 1A) except *DLK1*, which had normal expression in the adrenal gland, pituitary, ovary, hypothalamus, and testis with low to no expression in other normal tissues (Supplementary Fig. 1B). We next assessed tumor expression of *DLK1* using RNA-seq data from a cohort of ∼1000 adult patients with treatment refractory metastatic cancers^8^. We observed high *DLK1* expression in a subset of refractory cancers such as sarcomas, SCLC, germ cell tumors, and grade 2 neuroendocrine tumors (Fig. 1A). High *DLK1* expression has also been recently observed in pediatric neuroblastoma^9^. Strikingly, almost all adrenal cancers, i.e. ACC and pheochromocytoma/paraganglioma (PCPG), expressed high levels of *DLK1* (Fig. 1A), which we also observed in the TCGA PanCancer dataset^10^ (Fig. 1B). While ACC and PCPG are both rare tumors of the adrenal gland (ACC incidence of ∼0.5-2 cases per million people per year and PCPG incidence of 2-8 cases per million people per year^11^), we focused further analysis on ACC as it is an aggressive, highly malignant cancer with an overall poor prognosis (5-year survival 20-25%) with an urgent need for new treatment options^12^. *DLK1* was the most highly expressed Notch ligand with little to no expression of *DLL3* across multiple ACC cohorts^13–15^ including a new RNA-seq cohort generated from ACC metastases (n=50) at our institution (Fig. 1C, Supplementary Tables 1 and 2). To validate *DLK1* expression, we performed DLK1 IHC across our cohort of ACC metastatic tumors and found 97% (n=28/29) of ACC patients were DLK1^+^ (mean H-score 147) with H-scores ranging from 10 to 300 (Fig. 1D). Thus, our data demonstrate DLK1 as a potential new surface immunotherapeutic target in multiple malignancies, particularly ACC.

**Figure 1.**
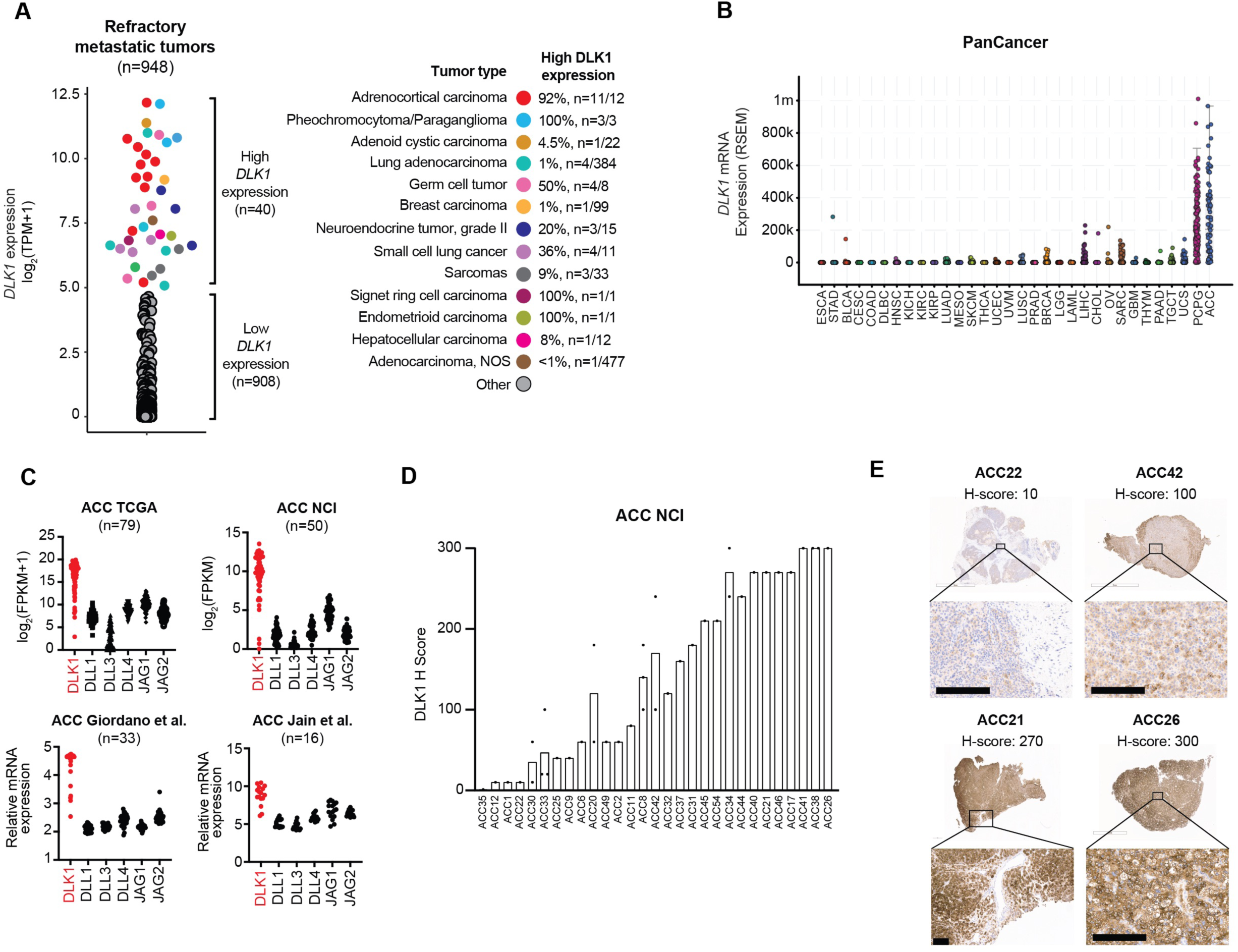
Identification of DLK1 as the most highly expressed Notch ligand in adrenocortical carcinoma. (**A**) *DLK1* mRNA expression across adult refractory metastatic cancers (n=948). Tumor types with high *DLK1* expression are highlighted in the colors shown. The percentage of each tumor type with high *DLK1* expression is shown on the right. (**B**) *DLK1* mRNA expression in the TCGA PanCancer dataset. (**C**) Expression of Notch ligands from four independent bulk ACC RNA-seq datasets. (**D**) Quantification of DLK1 IHC staining in ACC tumors with IHC images of four representation tumors with varying levels of DLK1 expression. Scale bars represent 200 μM. IHC: immunohistochemistry.

### ADCT-701, an antibody drug conjugate targeting DLK1, induces cytotoxicity in ACC through apoptosis and bystander killing

Given the high and near ubiquitous expression of DLK1 in ACC, we next sought to determine if DLK1 could be targeted in ACC using a DLK1-directed antibody-drug conjugate (ADCT-701). ADCT-701 consists of a humanized anti-DLK1 monoclonal IgG1 antibody coupled to SG3199, a pyrrolobenzodiazepine (PBD) dimer, which causes potent, cytotoxic DNA interstrand cross-linking of the minor groove of DNA^16^ (drug-to-antibody ratio∼1.8) via a Val-Ala cleavable linker and HydraSpace^TM^ utilizing the GlycoConnect^TM^ technology (Fig. 2A). We determined the cytotoxicity of ADCT-701 *in vitro* using three established ACC cell lines with varying levels of DLK1 surface expression (Fig. 2B). Compared to the isotype-control ADC (B12-PL1601), ADCT-701 inhibited cell growth in DLK1^+^ CU-ACC1 and H295R cells, but not in DLK1^-^ CU-ACC2 cells (Fig. 2C). However, DLK1^-^ CU-ACC2 cells, similar to CU-ACC1 and H295R cells, were sensitive to the PBD payload of ADCT-701 (Supplementary Fig. 2A). We next used CRISPR-Cas9 gene editing in the CU-ACC1 cell line and established multiple single cell clones with complete loss of DLK1 (Supplementary Fig. 2B, C). In several CU-ACC1 DLK1 KO clones, ADCT-701 cytotoxicity was abrogated (Supplementary Fig. 2D) thereby validating the DLK1-specific cytotoxicity of ADCT-701. We observed similar findings in the DLK1^-^ PCPG cell line hPheo1^17^ (Supplementary Fig. 2E, F). Taken together, these data demonstrate that ADCT-701 exhibits *in vitro* cytotoxic activity in ACC in a DLK1-dependent manner.

**Figure 2.**
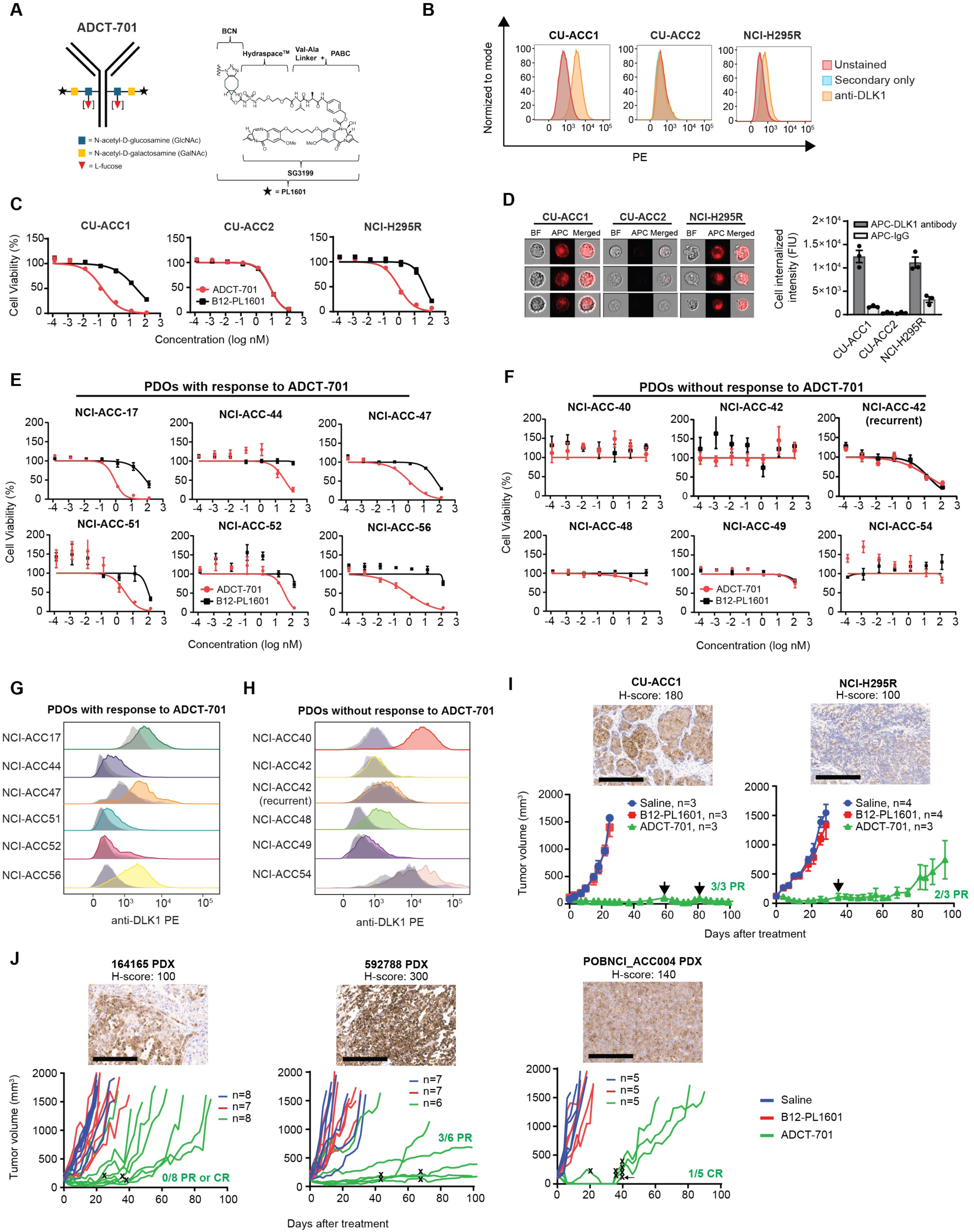
ADCT-701, a DLK1 targeting antibody-drug conjugate, has potent *in vitro* activity and induces robust *in vivo* anti-tumor responses in adrenocortical carcinoma. (**A**) Schematic structure of ADCT-701, a DLK1 targeting antibody drug conjugate. (**B**) Representative surface expression of DLK1 among ACC cell lines: CU-ACC1, CU-ACC2, and H295R. (**C**) ADCT-701 cytotoxicity among CU-ACC1, CU-ACC2, and H295R cells. Cells were treated with ADCT-701 and B12-PL1601 (non-targeted control ADC) for 7 days. Each point represents the mean±SEM. (**D**) Representative imaging flow cytometry images and signal intensity analysis (n=3 biological replicates) showing cellular internalization of DLK1 antibodies in CU-ACC1, CU-ACC2, and H295R. (**E**) Cytotoxic activity of ADCT-701 responsive (n=6) and (**F**) non-responsive (n=6) ACC patient-derived organoids (PDOs). Flow cytometry histograms assessing DLK1 among ADCT-701 (**G**) responsive and (**H**) non-responsive ACC PDOs. Shaded gray histograms represent unstained controls for each condition. (**I**) CU-ACC1 and H295R xenograft tumor growth curves after treatment with saline, B12-PL1601, or ADCT-701 (1 mg/kg). Additional doses of ADCT-701 indicated by arrows. (**J**) ACC PDXs 164165, 592788, and POBNCI_ACC004 tumor growth curves after treatment with saline, B12-PL1601 or ADCT-701 (1 mg/kg). X symbols indicate the administration of ADCT-701 re-dosing. Arrow indicates unexpected death of 1 POBNCI_ACC004 tumor-bearing mouse prior to endpoint. DLK1 immunohistochemistry with H-scores shown above each individual xenograft or PDX tumor. Scale bars represent 200 μM.

We next evaluated the mechanism by which ADCT-701 induces cell death. Cellular internalization of ADC after binding to the surface target is an essential step for ADC cytotoxicity^18^. To demonstrate that our anti-DLK1 mAb could be internalized efficiently, we quantified the cellular internalization rates of DLK1 antibody across ACC cell lines with varying DLK1 expression levels using imaging flow cytometry. DLK1 antibody was rapidly internalized in DLK1^+^ CU-ACC1 and H295R cells but not in DLK1^-^ CU-ACC2 cells (Fig. 2D). Next, consistent with previous studies demonstrating PBD induced DNA interstrand crosslinks results in cell cycle arrest^19^, we found that CU-ACC1 and H295R cells treated with ADCT-701 were blocked in the G2/M phase (Supplementary Fig. 3A and Supplementary Fig. 4A). We then determined the presence of DNA double-strand breaks by γH2AX, as well as apoptosis by cleaved caspase-3 and cleaved poly (adenosine diphosphate-ribose) polymerase (PARP). γH2AX, cleaved caspase 3, and cleaved PARP were upregulated in CU-ACC1 and H295R cells after ADCT-701 but not after B12-PL1601 treatment (Supplementary Fig. 3B) and ADCT-701 significantly increased apoptosis (Annexin V+/PI+) compared to untreated and B12-PL1601 treated cells (Supplementary Fig. 3C and Supplementary Fig. 4B). Collectively, these results suggest that ADCT-701 treatment leads to DNA double-strand breaks, G2/M arrest, and ultimately apoptosis.

Due to the heterogeneous expression of DLK1 in ACC (Fig. 1E), we next assessed for potential bystander killing^18^ using a system in which DLK1 KO CU-ACC1 cells were cultured with DLK1-expressing parental CU-ACC1 cells at various ratios. We observed greater cytotoxicity of DLK1 KO CU-ACC1 cells than expected with no cytotoxicity observed with B12-PL1601 treatment (Supplementary Fig. 3D). We further investigated bystander killing by conditioned media transfer experiments in which DLK1^+^ CU-ACC1 cells were treated with ADCT-701 or B12-PL1601 before transferring the media to DLK1^+^ CU-ACC1 cells or DLK1 KO CU-ACC1 cells. ADCT-701 conditioned media induced cytotoxicity in DLK1^+^ CU-ACC1 cells similar to treatment with ADCT-701 (Supplementary Fig. 3E). ADCT-701 conditioned media also elicited bystander killing, as demonstrated by greater cytotoxicity in DLK1 knockout CU-ACC1 cells compared with B12-PL1601 conditioned media or ADCT-701 treatment (Supplementary Fig. 3E). Overall, these results indicate that ADCT-701 not only can target DLK1^+^ cells but can also indirectly induce cytotoxicity in DLK1^-^ cells.

### ADCT-701 has potent in vitro activity in DLK1^+^ ACC patient-derived organoids and induces robust anti-tumor responses in ACC cell line-derived and patient-derived xenografts

Since ACC is a rare cancer type with few available human cell lines^20^, we sought to validate the *in vitro* cytotoxicity of ADCT-701 in a newly developed cohort of ACC short-term patient-derived organoids (PDOs) (defined as less than 5 total passages) (Supplementary Table 3). Overall, we found 50% (n=6/12) of PDOs responded to ADCT-701 (Fig. 2E) and 50% (n=6/12) of PDOs had no response (Fig. 2F). As expected, all ADCT-701 responders were DLK1^+^ (Fig. 2G), However, among ADCT-701 non-responders, 50% (n=3/6) were still DLK1^+^ (Fig. 2H) suggesting that ADCT-701 sensitivity is influenced by factors other than DLK1 expression.

Next, to further explore the potential for targeting DLK1 in ACC, we evaluated responses to ADCT-701 among DLK1^+^ human ACC cell line-derived xenograft and ACC patient-derived (PDX) models (Fig. 2I, J). ADCT-701 treatment elicited durable anti-tumor responses and significantly prolonged the survival of both CU-ACC1 and H295R tumor-bearing mice compared with tumor-bearing mice treated with saline or B12-PL1601 (Fig. 2I and Supplementary Fig. 5A). However, H295R tumors eventually became resistant to ADCT-701, whereas CU-ACC1 tumors remained sensitive to ADCT-701 for up to 100 days. While the DLK1 antibody within ADCT-701 targets human not mouse DLK1, no body weight loss in mice was observed with ADCT-701 treatment (Supplementary Fig. 5B) suggesting minimal off-target payload activity.

We next investigated the anti-tumor activity of ADCT-701 among three DLK1^+^ ACC PDX models: 164165, 592788, and POBNCI_ACC004 (Fig. 2J and Supplementary Fig. 6A). All three models were validated as ACC tumors based on IHC expression of the common neuroendocrine marker synaptophysin, the adrenal specific marker SF1, and the cell proliferation marker Ki67 (Supplementary Fig. 6B-D). ADCT-701 treatment led to tumor growth inhibition and significant lengthening of survival among 164165 PDX and 592788 PDX mice (Fig. 2J and Supplementary Fig. 5C). Strikingly, ADCT-701 induced complete responses in all treated POBNCI_ACC004 PDX tumors (5/5). However, 3/5 POBNCI_ACC004 PDX tumors quickly recurred and did not respond to ADCT-701 re-treatment (Fig. 2J). Similar to treated xenografts, ADCT-701 was well-tolerated in PDX models as determined by body weight measurements (Supplementary Fig. 5D).

### ABCB1, a drug efflux protein, mediates intrinsic and acquired ADC and chemotherapy resistance among DLK1^+^ ACC pre-clinical models

We next sought to decipher mechanisms of resistance to ADCT-701 in our DLK1^+^ ACC pre-clinical models. As payload insensitivity is known to mediate ADC resistance^18^, we first tested the cytotoxicity of PBD across DLK1^+^ ADCT-701 responder and non-responder PDOs. Strikingly, we observed extreme resistance to PBD among non-responder PDOs, with PBD average IC50s of 97 nM in NCI-ACC40, 31 nM in NCI-ACC48, and 54 nM in NCI-ACC54, which represents close to 1000x greater resistance than previously reported for PBD^21^ (Fig. 3A). We then explored the activity of common therapies used to treat ACC in the clinic^12^ (mitotane, etoposide, doxorubicin, and carboplatin) among two DLK1^+^ ACC PDOs with and without response to ADCT-701 and PBD (NCI-ACC51 and NCI-ACC48, respectively) that were able to grow in culture longer than other PDOs (greater than 5 passages). We observed resistance to chemotherapy in the NCI-ACC48 PDO compared to the NCI-ACC51 PDO but no difference in mitotane activity (Supplementary Fig. 7A). We next looked for potential mechanisms of resistance to PBD by analyzing our NCI-ACC RNA-seq patient dataset for expression of drug transporter genes previously implicated in resistance to PBD-ADCs^22^ as well as commonly upregulated drug efflux pumps of the ABC transporter family^21^. *ABCB1* (MDR1 or p-glycoprotein) and *ABCG2* had the greatest difference in expression between NCI-ACC48 and NCI-ACC51 patient tumors (Fig. 3B) suggesting these drug efflux genes could explain the difference in PBD resistance in corresponding PDOs. Indeed, the NCI-ACC48 PDO had much higher surface expression of ABCB1 than the NCI-ACC51 PDO (Fig. 3C). Furthermore, co-treatment of 3 different ABCB1 inhibitors (valspodar, elacridar, and tariquidar) with PBD and ADCT-701 all showed dramatic reversal of resistance in the NCI-ACC48 PDO (Fig. 3D and Supplementary Fig. 7B). ABCB1 inhibitors more modestly increased sensitivity to ADCT-701 and PBD in the NCI-ACC51 PDO demonstrating a functionally lower level of ABCB1 activity in this model compared to the NCI-ACC48 PDO (Supplementary Fig. 7C). Thus, these results indicate that primary *in vitro* resistance to ADCT-701 can be mediated by high expression and activity of the drug efflux protein ABCB1.

**Figure 3.**
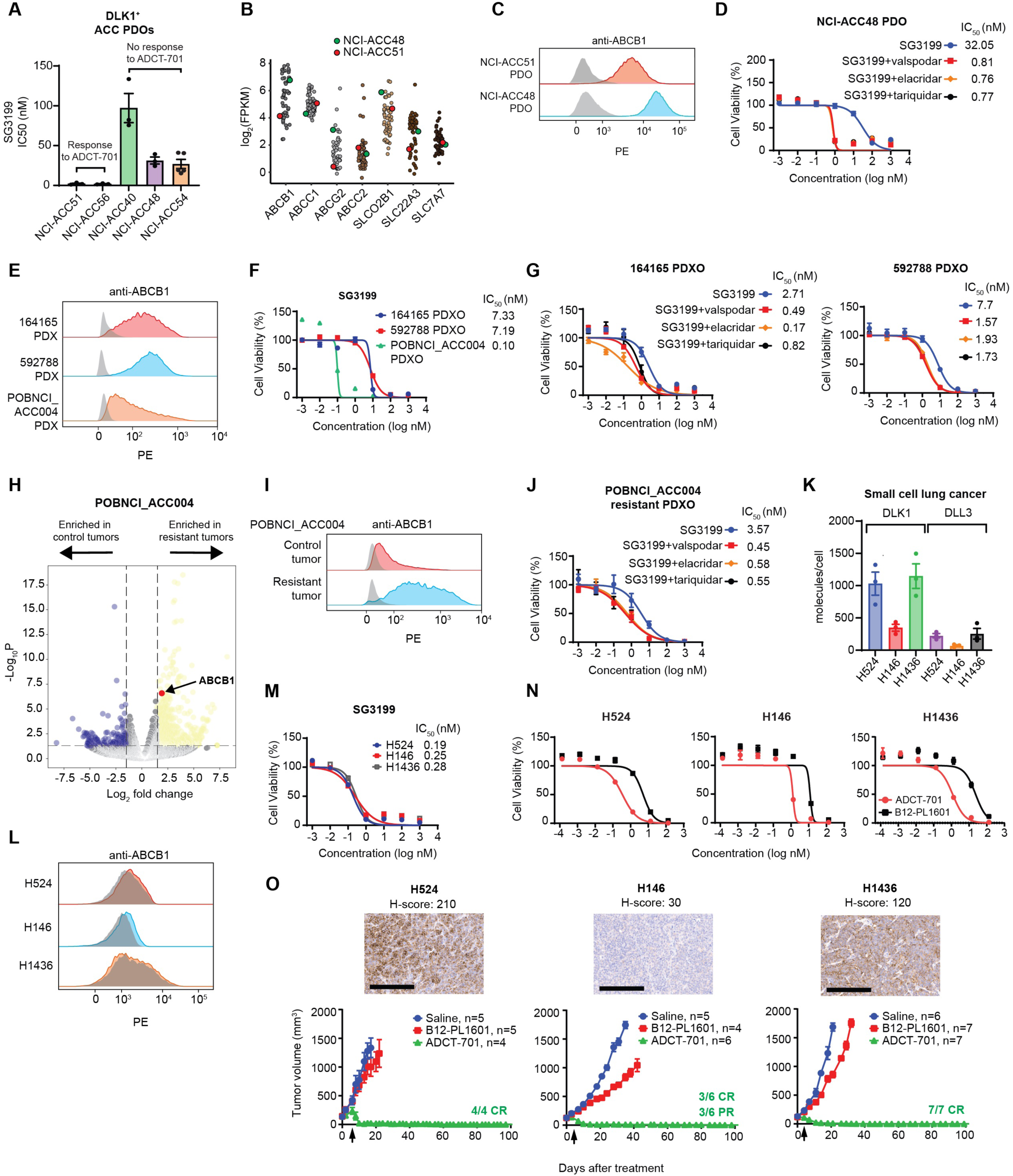
ABCB1, a drug efflux protein, mediates intrinsic and acquired resistance to ADCT-701. (**A**) Cytotoxicity of SG3199 among ADCT-701 responsive and non-responsive DLK1^+^ ACC patient-derived organoids (PDOs). (**B**) Drug transporter mRNA expression in NCI-ACC patients as measured by RNA-seq. (**C**) Flow cytometry histograms assessing ABCB1 of DLK1^+^ ADCT-701 responsive (NCI-ACC51) and non-responsive (NCI-ACC48) ACC PDOs. (**D**) SG3199 cytotoxicity in the NCI-ACC48 PDO with and without treatment with ABCB1 inhibitors (1 μM valspodar, 10 μM elacridar and 1 μM tariquidar). (**E**) Flow cytometry histograms assessing ABCB1 among 164165, 592788 and POBNCI_ACC004 PDXs. (**F**) SG3199 cytotoxicity in 164165, 592788 and POBNCI_ACC004 PDX-derived organoids. (**G**) SG3199 cytotoxicity in the 164165 and 592788 PDX-derived organoids treated with or without ABCB1 inhibitors (1 μM valspodar, 10 μM elacridar and 1 μM tariquidar). (**H**) Volcano plot of differentially expressed genes in control tumors versus post-ADCT-701 acquired resistant tumors in POBNCI_ACC004 PDX. (**I**) Flow cytometry histograms assessing ABCB1 among ADCT-701 resistant and control POBNCI_ACC004 PDX tumors. (**J**) SG3199 cytotoxicity in the ADCT-701 resistant POBNCI_ACC004 PDX-derived organoid treated with or without ABCB1 inhibitors (1 μM valspodar, 10 μM elacridar and 1 μM tariquidar). **(K)** DLK1 molecules/cell relative to DLL3 among small cell lung cancer (SCLC) cell lines (H524, H146, and H1436). (**L**) Flow cytometry histograms assessing ABCB1 in SCLC cell lines. **(M)** SG3199 cytotoxicity in SCLC cell lines. Cells were treated with SG3199 for 3 days. **(N)** ADCT-701 cytotoxicity among SCLC cell lines. Cells were treated with ADCT-701 or B12-PL1601 for 7 days. Each point represents the mean±SEM. **(O)** Tumor growth curves of SCLC xenograft models after treatment with saline, B12-PL1601 or ADCT-701 (1 mg/kg) treatment. Arrows indicate re-treatment with ADCT-701. Scale bars represent 200 μM. For flow cytometry histograms, shaded gray histograms represent isotype controls for each condition.

We next sought to determine if ABCB1 expression and activity could also explain differences in ADCT-701 *in vivo* activity across our ACC cell line-derived xenograft and PDX models. Among our ACC xenografts, CU-ACC1 had lower surface ABCB1 expression than H295R (Supplementary Fig. 8A), which could at least partially explain the long-term tumor control with ADCT-701 treatment in CU-ACC1 but not H295R tumors (Fig. 2I). Among our ACC PDXs, there was considerably lower surface ABCB1 expression in POBNCI_ACC004 (Fig. 3E), which had initial complete responses to ADCT-701, compared to both 164165 and 592788 (Fig. 3E), which had partial but no complete anti-tumor responses to ADCT-701. To further test the role of ABCB1 in relation to ADCT-701 activity in these models, we developed PDX-derived organoids from untreated POBNCI_ACC004, 164165 and 592788 PDX tumors (i.e. PDXOs). Consistent with our *in vivo* data, POBNCI_ACC004 PDXO was much more sensitive to SG3199 than 164165 and 592788 PDXOs (Fig. 3F). ABCB1 inhibitors also increased sensitivity to SG3199 among 164165 and 592788 PDXOs (Fig. 3G) demonstrating that ABCB1 drug efflux activity is a mechanism of intrinsic resistance to ADCT-701 in these two models.

Although POBNCI_ACC004 PDX tumors had initial complete responses to ADCT-701 treatment, these tumors quickly relapsed and were unresponsive to additional ADCT-701 doses (Fig. 2J). Therefore, to assess mechanisms of acquired resistance to ADCT-701, we performed RNA-seq on POBNCI_ACC004 PDX tumors resistant to ADCT-701 treatment (n=3) and saline treated control tumors (n=4). Using differential gene expression analysis, we observed upregulation of *ABCB1* expression in POBNCI_ACC004 PDX tumors resistant to ADCT-701 compared to controls (Fig. 3H and Supplementary Table 4). Surface ABCB1 expression was also highly upregulated in an POBNCI_ACC004 resistant compared to an untreated POBNCI_ACC004 control tumor (Fig. 3I). We then developed a PDXO from a POBNCI_ACC004 resistant tumor and found that ABCB1 inhibitors re-sensitized the POBNCI_ACC004 resistant PDXO to PBD and ADCT-701 (Fig. 3J and Supplementary Fig. 8C) demonstrating the role of this drug efflux transporter in mediating acquired resistance to ADCT-701. Lastly, unlike in neuroblastoma^23^, we found no difference in DLK1 expression by IHC between pre- and post-ADCT-701 treated ACC PDX tumors (Supplementary Fig. 8D) suggesting that selection and outgrowth of DLK1 negative cells does not contribute to ADCT-701 acquired resistance in ACC.

### ADCT-701 elicits complete, durable responses in DLK1^+^ small cell lung cancer tumors without ABCB1 expression

As we found DLK1 to be expressed in a subset of metastatic cancers apart from ACC (Fig. 1A), we hypothesized that ADCT-701 would be highly effective against DLK1^+^ tumors with low or no ABCB1 expression such as SCLC. We therefore screened SCLC cell lines for expression of DLK1 and found 22% (n=11/51) were DLK1^+^ (Supplementary Fig. 9A). We then selected three DLK1^+^ SCLC cell lines (H524, H146, and H1436) and confirmed that cell surface DLK1 expression was at a level equal to or higher than the known SCLC target DLL3 (Fig. 3K and Supplementary Fig. 9B). All three SCLC cell lines also lacked surface ABCB1 expression (Fig. 3L) and were highly sensitive to both SG3199 and ADCT-701 *in vitro* (Fig. 3M, N). *In vivo*, ADCT-701 treatment resulted in complete responses and long-term tumor-free survival compared to B12-PL1601 and saline in all three of the SCLC xenograft models (Fig. 3O and Supplementary Fig. 9C) without appreciable body weight loss (Supplementary Fig. 9D). Notably, the SCLC H146 xenograft, which had very low DLK1 expression (H-score 30), also had long-term complete responses with ADCT-701 treatment (Fig. 3O). Thus, our results indicate that ADCT-701 can effectively target DLK1^+^ tumors with low or no ABCB1 expression.

### DLK1 is a major regulator of ABCB1, adrenocortical differentiation, and chemoresistance in ACC

We next sought to assess whether DLK1 has a functional role in ACC using our established CU-ACC1 DLK1 KO cells (Supplementary Fig. 2B, C). As DLK1 has been shown to activate or inhibit NOTCH1 signaling in several model systems^24^, we assessed expression of NOTCH1 in DLK1 KO compared to DLK1 WT parental cells and observed upregulation of total NOTCH1 and the active, intracellular domain (ICD) of NOTCH1 (N1ICD) (Fig. 4A). There was also a dramatic reduction in the NE protein synaptophysin (Fig. 4A) after DLK1 KO consistent with the known role of active Notch signaling in downregulating NE gene expression^25^. Correspondingly, we observed a significant negative correlation between *NOTCH1* and *DLK1* expression across TCGA ACC tumors (Fig. 4B). *DLK1* and *NOTCH1* expression were also significantly higher and lower, respectively, in ACC tumors compared to the normal adrenal gland (Fig. 4C).

**Figure 4.**
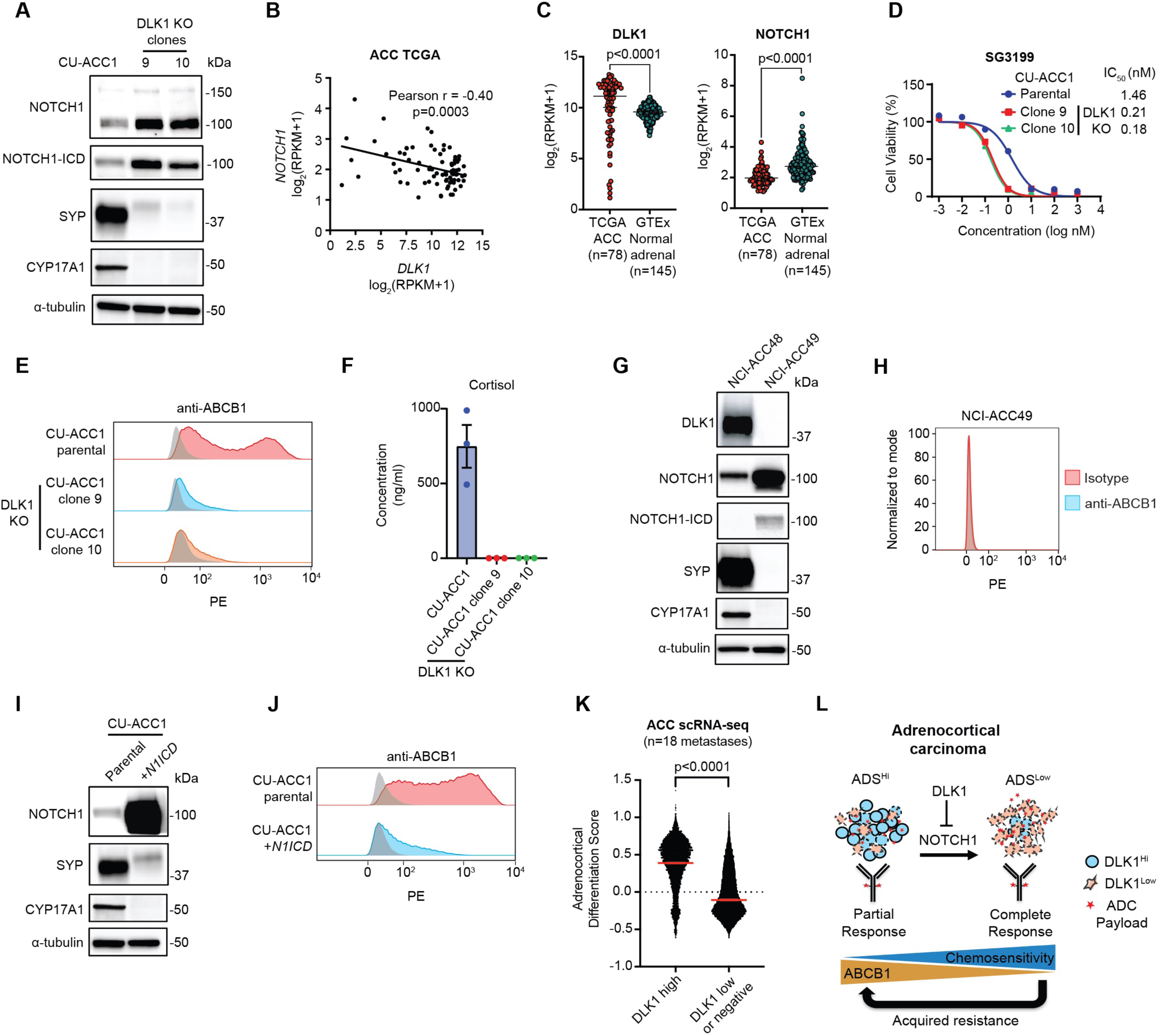
DLK1 is a major regulator of ABCB1, adrenocortical differentiation, and chemoresistance in adrenocortical carcinoma. **(A)** Immunoblot analysis of NOTCH1 signaling, total NOTCH1 and NOTCH1 intracellular domain (ICD), NE marker synaptophysin (SYP), and loading control (α-tubulin) proteins with and without DLK1 KO in CU-ACC1 cells. Two single-cell KO clones are shown. **(B)** Correlation between *NOTCH1* and *DLK1* expression among TCGA ACC tumors. **(C)** *DLK1* and *NOTCH1* expression in TCGA ACC tumors and normal adrenals. **(D)** SG3199 cytotoxicity in CU-ACC1 parental and DLK1 KO clones. **(E)** Flow cytometry histograms assessing ABCB1 in CU-ACC1 cells with and without DLK1 KO. **(F)** Concentration of cortisol in conditioned media from CU-ACC1 parental and DLK1 KO clones. **(G)** Immunoblot analysis of DLK1, total NOTCH1 and NOTCH1-ICD, SYP, the steroidogenic enzyme CYP17A1, and α-tubulin proteins in DLK1^+^ NCI-ACC48 and DLK1^-^ ACC49 patient-derived organoids. **(H)** Flow cytometry histograms assessing ABCB1 in DLK1 negative NCI-ACC49 patient-derived organoids. **(I)** Immunoblot analysis of DLK1, total NOTCH1 (to detect the NOTCH1-ICD plasmid expression), SYP, CYP17A1, and α-tubulin proteins in CU-ACC1 cells with and without NOTCH1-ICD overexpression. **(J)** Flow cytometry histograms assessing ABCB1 in CU-ACC1 cells with and without *N1ICD* overexpression. **(K)** Single cell RNA-seq adrenocortical differentiation score comparing DLK1 high cells to DLK1 low or negative cells from 18 ACC metastatic tumors. **(L)** Model summarizing the findings of the current study. For flow cytometry histograms, shaded gray histograms represent isotype controls for each condition.

Given that prior work has shown Notch-active tumors with low NE gene expression (i.e. non-NE) to be chemoresistant^26^, we performed cytotoxicity assays with PBD in CU-ACC1 cells with and without DLK1 KO. Surprisingly, DLK1 KO cells were much more sensitive to PBD (Fig. 4D) and chemotherapeutics such as etoposide and doxorubicin (Supplementary Fig. 10A). However, DLK1 KO cells displayed no change in proliferation compared to parental cells. (Supplementary Fig. 10B). Strikingly, DLK1 KO had near complete loss of ABCB1 surface expression compared to parental cells, which showed a broad, bi-modal distribution of ABCB1 (Fig. 4E). In DLK1 KO cells, we also observed strong downregulation of steroidogenesis protein CYP17A1 (Fig. 4A), which is highly expressed in the adrenal cortex and composes part of the Adrenocortical Differentiation Score (ADS)^14^. However, CYP17A1 expression was not decreased with siRNA downregulation of DLK1 (Supplementary Fig. 10C) suggesting longer-term NOTCH1 signaling, known to be required to induce transdifferentiation^26^, is required to induce changes in adrenocortical differentiation. CU-ACC1 DLK1 KO clones also had dramatically lower secretion of cortisol compared to parental cells (Fig. 4F). As we observed DLK1 KO cells to be completely adherent compared to a mixed phenotype (suspension and adherent) of CU-ACC1 parental cells (Supplementary Fig. 10D) we isolated suspension and adherent CU-ACC1 cells (Supplementary Fig. 10E). Suspension CU-ACC1 cells showed high expression of DLK1 and SYP and low expression of N1ICD (Supplementary Fig. 10F). In contrast, adherent CU-ACC1 cells showed low expression of DLK1 and SYP and high expression of N1ICD (Supplementary Fig. 10F). CU-ACC1 adherent cells also had lower expression of surface ABCB1 with modest differences in chemosensitivity compared to CU-ACC1 suspension cells (Supplementary Fig. 10G, H).

To validate our DLK1 KO results, we assessed expression of NOTCH1, NE and adrenocortical differentiation proteins in an ACC PDO with high DLK1 expression (NCI-ACC48) and an ACC PDO with no DLK1 expression (NCI-ACC49) (Figure 4G). We observed higher expression of N1ICD and much lower expression of SYP and CYP17A1 in the DLK1^-^ NCI-ACC49 PDO compared to the DLK1^+^ NCI-ACC48 PDO (Figure 4G). The NCI-ACC49 PDO was also highly sensitive to SG3199, etoposide and doxorubicin (Supplementary Fig. 10I) and ABCB1 was not expressed (Fig. 4H). We next overexpressed *N1ICD* in CU-ACC1 cells, and similar to DLK1 KO cells, we observed decreased expression of synaptophysin and near complete downregulation of CYP17A1 and ABCB1 (Figure 4I, J). However, there were only minor differences in chemosensitivity between CU-ACC1 and *N1ICD*-overexpressing CU-ACC1 cells (Supplementary Fig. 10J) suggesting DLK1 KO may mediate chemosensitivity through additional mechanisms apart from ABCB1 expression and NOTCH1 signaling. Lastly, we analyzed single-cell RNA-seq data from 18 ACC metastatic tumors (*Aber et al. manuscript in submission*) and observed a significantly higher ADS among cells with high *DLK1* expression compared to cells with low or no *DLK1* expression (Fig. 4K) supporting our experimental results. Altogether, our data suggest a model by which DLK1, through inhibition of NOTCH1 signaling, maintains adrenocortical differentiation and high ABCB1 expression and imparts ADC and chemoresistance in ACC (Fig. 4L). Based on our data, DLK1-directed ADCs would also be expected to have greater activity in ACC tumors with positive but low DLK1 expression due to decreased adrenocortical differentiation and ABCB1 expression (Fig. 4L).

### A first-in-human phase I clinical trial of ADCT-701 in patients with ACC and neuroendocrine neoplasms

Based on the pre-clinical efficacy of ADCT-701 in ACC and SCLC, as well as parallel data in neuroblastoma^9^, a first-in-human phase 1 clinical trial was developed to test the safety and preliminary efficacy of ADCT-701 in adult patients with ACC and neuroendocrine neoplasms. This trial (NCT06041516) is currently recruiting patients with the primary objective to determine the maximum tolerated dose (MTD) of ADCT-701.

## Discussion

In this work, we have identified DLK1 as a cancer cell surface antigen that can be successfully targeted with an ADC in pre-clinical models of refractory metastatic cancers, namely ACC and SCLC. While ADCs are an effective and increasingly common cancer therapy, approval is currently limited to select malignancies (i.e., breast cancer, urothelial cancers, and ovarian cancers) with overall few antigen targets (i.e. TROP2, nectin-4, HER2, tissue factor, and folate receptor alpha^18^). Thus, identifying new and optimal cell surface targets, such as DLK1, is an important step towards broadening the therapeutic potential of ADCs. Indeed, based on our pre-clinical data, we have initiated a first-in-human phase 1 clinical trial with an ADC targeting DLK1 (NCT06041516). To our knowledge, this is the first ADC clinical trial for patients with ACC and neuroendocrine neoplasms including rare neuroendocrine malignancies such as PCPG and adult neuroblastoma.

In addition to identifying DLK1 as a new immunotherapeutic target, we demonstrate a novel role for DLK1 in conferring ADC and chemoresistance through high expression and activity of the multidrug efflux pump ABCB1. Intrinsic resistance to ADCs is common in the clinic with the ABCB1 drug transporter considered to be one potential mechanism^27^. Specifically in ACC, chemotherapy resistance has long been attributed to high expression of ABCB1^28,29^, which is known to be one of several genes highly expressed in the adrenal cortex^14^. Our findings could thus have important implications for ACC treatment as inhibition of DLK1 could be a strategy to reduce resistance to chemotherapy in ACC, particularly the EDP (etoposide, doxorubicin, cisplatin) regimen which includes two chemotherapeutic drugs known to be ABCB1 transport substrates (etoposide and doxorubicin). EDP chemotherapy is widely used to treat advanced ACC but has not improved overall survival^1,30^.

A key new mechanistic finding from this work is the identification of DLK1 as a driver of adrenocortical. Previous work has identified several key regulators of adrenocortical differentiation such as the histone methyltransferase EZH2^31,32^ ^9^ and the deSUMOylating enzyme SENP2^33,34^. Our data suggest that DLK1 maintains adrenal steroidogenic cell differentiation, which is concordant with recent data from a spatial transcriptomic analysis of DLK1^+^ and DLK1^-^ ACC tumor regions^35^. Moreover, our data are consistent with the known role of DLK1 regulating cellular differentiation processes such as adipogenesis, hematopoiesis, stem cell homeostasis, neurogenesis, angiogenesis, and muscle regeneration^36^. In regard to ABCB1, we demonstrate that long-term NOTCH1 activation and de-differentiation of ACC are required to downregulate ABCB1. As prior work has demonstrated NOTCH1 as a positive regulator of ABCB1 expression in several other solid tumor models^37^, our data highlights the context dependent nature of Notch signaling^30^.

Our experimental data also uncovers a role for DLK1 in transdifferentiating cells from a NE to non-NE state, which to our knowledge, has not been previously known, although DLK1 has been observed to be upregulated in NE tumors^31,32^. Interestingly, adrenocortical carcinomas are not typically categorized as NE tumors as they are not of neuroepithelial origin and they generally lack expression of NE genes such as chromogranin A^38^. Indeed, neuroendocrine scoring systems have used gene expression data from the normal adrenal cortex to identify non-NE genes compared to NE genes in the normal adrenal medulla^39^. However, ACC tumors commonly express NE genes such as synaptophysin^40^ and our data demonstrate that synaptophysin is regulated by DLK1 through NOTCH1. Low or no expression of synaptophysin on routine ACC tumor specimens, albeit likely low in prevalence, could indicate a non-NE, less adrenocortical differentiated state (i.e. DLK1^low^/NOTCH1^high^/ABCB1^low^) with sensitivity to chemotherapy or an ADC. Counterintuitively, our data suggest that high DLK1 expression may not be an optimal biomarker for a DLK1-directed ADC as DLK1^high^ tumors would be expected to have high ABCB1 expression and thereby demonstrate payload resistance. Rather, tumors with low DLK1 expression (which are able to bind and internalize a DLK1-directed ADC) may exhibit the most optimal ADC response due to low ABCB1 expression.

The direct link we propose between DLK1, NOTCH1 signaling, and ABCB1 expression also suggest that ABCB1 inhibition could improve anti-tumor responses to DLK1-directed ADCs. However, a clinical trial testing the addition of the ABCB1 antagonist, tariquidar, to chemotherapy among ACC patients was stopped prematurely due to toxicity^41^. Off-target toxicity of the combination of ABCB1 inhibitors and chemotherapy (such as to bone marrow whose stem cells are protected from chemotherapy by expression of ABC efflux transporters such as ABCG and ABCB1) should be much less of a concern when ABCB1 inhibitors are combined with targeted therapy such as ADCs, which have reduced off-target toxicity. One other strategy for future ADCs could be to use payloads that are not substrates for drug efflux transporters.

Beyond ADCs, degrader-antibody conjugates^18^ targeting DLK1 may be a strategy to downregulate DLK1, which could potentially sensitize ACC tumors to chemotherapy or other ADCs. Moreover, there are now multiple immunotherapeutic strategies to target cancer cell surface proteins such as CAR T cells and bi-specific T cell engagers (BiTEs). Indeed, DLK1-directed CAR T cells have been shown to have pre-clinical efficacy among DLK1-expressing hepatocellular carcinoma models^42^. CAR T cells may be a particularly attractive option in ACC given the high level of chemoresistance; however, CAR T cells generally have more toxicity than ADCs^43^ and thus it may advisable to accrue safety information on targeting DLK1 from our phase 1 study before pursuing clinical testing of CAR T cells. Although DLK1 has minimal expression across most normal tissues, there is high expression in several organs such as the adrenal gland, particularly the adrenal medulla compared to the adrenal cortex^9,44^. ADCT-701 treatment may thus lead to adrenal hormone deficiency requiring supplementation with mineralocorticoids and/or corticosteroids. Another active phase 1 trial targeting DLK1 with an ADC in advanced cancers (NCT06005740)^45^ using monomethyl auristatin E (MMAE) as the payload (also an ABCB1 substrate), may also provide additional safety information.

There are several limitations to our study. While we demonstrate that DLK1 is a regulator of ABCB1 expression and thereby sensitivity to a DLK1-directed ADC and chemotherapy, there are likely additional variables which affect ADC and chemotherapy sensitivity that we are unable to account for in this study. Moreover, while we focused on the role of DLK1 in mediating ADC resistance in our functional studies, DLK1 is known to regulate cancer stemness^36,46,47^ and tumor progression^48^. Thus, further investigation into the role of DLK1 in ACC tumorigenesis will be important. Also, although our study focused on ACC and SCLC, our screening data revealed high *DLK1* expression across several additional metastatic tumor types such as germ cell tumors and sarcomas and recent parallel work has demonstrated DLK1 as an immunotherapeutic cell surface target in pediatric neuroblastoma^9^. Thus, a biomarker-based assessment of DLK1 across a broader group of malignancies could be a future clinical approach for DLK1-directed immunotherapeutic clinical trials.

In summary, we have identified DLK1 as a new immunotherapeutic target in ACC and neuroendocrine neoplasms such as SCLC. We have also demonstrated DLK1 as an important driver of chemotherapy and ADC resistance through regulation of the drug efflux pump ABCB1. Our data support the clinical testing of targeting DLK1 with an ADC in ACC and other neuroendocrine neoplasms and identify DLK1 as an important cell surface target for future immunotherapeutic approaches.

## Materials and Methods

### ACC patient tumor specimens

ACC patient tumors were collected from surgical resection of metastatic sites at the NIH Clinical Center under NIH Institutional Review Board protocols (NCT05237934, NCT01109394, and NCT03739827). Tumor tissue from surgical resections was used for organoid experiments and DLK1 IHC. RNA-sequencing was performed from formalin-fixed tissue acquired either from surgical resected tissue or from archival tissue. A summary of ACC patient tumors with associated RNA-seq, DLK1 IHC and/or organoid assay data used in this study are shown in Supplementary Table 1.

### Bulk and single-cell RNA sequencing

Formalin-fixed, paraffin-embedded (FFPE) tumor tissue samples were prepared for bulk RNA sequencing (RNA-Seq). RNA-seq libraries were prepared using Illumina TruSeq RNA Access Library Prep Kit or Total RNA Library Prep Kit according to the manufacturer’s protocol (Illumina). The NCI CCBR RNA-seq pipeline (https://github.com/skchronicles/RNA-seek.git) was used for further processing. In summary, STAR (2.7.6a) was run to map reads to hg38 (release 36) reference genome. Then, RSEM was used to generate gene expression values in log_2_(FPKM+1). We applied the "RemoveBatchEffect" function from the package Limma to remove the impact of the library preparation protocols (access or totalRNA). Single cell RNA-seq was performed as described in *Aber et al. manuscript in submission*. In brief, single cell suspensions from 18 ACC liver and/or lung metastases were sequenced on the 10x Genomics Chromium Platform targeting 6,000 cells per sample. Data was processed using the cellranger pipeline and downstream analysis performed in R using Seurat. Our analysis categorized *DLK1* expression as high or low/negative in malignant cells only (identified by copy number variation using inferCNV). An adrenocortical differentiation score was calculated using genes from a previously published adrenocortical differentiation gene set apart from DLK1, which was excluded^14^.

### Tumor cells isolation and organoid culture

To generate organoid cultures, fresh ACC patient tumors were minced into tiny fragments in the sterile dish. Tumor fragments were performed to enzymatic digestion in advanced DMEM/F12 supplemented with 1x Glutamax and 10 mM HEPS buffer containing collagenase type IV (200 U/ml, Sigma–Aldrich) and DNase I (50 U/ml, Sigma–Aldrich) on an orbital shaker for 1 hr at 37 °C and filtered through 70 μm strainers. The mixture was spun for 5 min at 1500 rpm. The cell pellet was treated with 1 x RBC lysis buffer (Sigma-Aldrich #) for 5 min at room temperature to remove the red blood cells and then spun for 5 min at 1500 rpm. Single cell suspensions were seeded on a Matrigel dome. Matrigel was mixed with tumor cells in minimum basal medium (MBM) consisting of 1x N2 supplement (Thermo Fisher Scientific #17502048), 1x B27 supplement (Thermo Fisher Scientific #17504044), 50 ng/mL EGF (Thermo Fisher Scientific #PHG0311), 20 ng/mL bFGF (STEMCELL Technologies #78003), 100 ng/mL IGF-2 (STEMCELL Technologies #78023), and 10 μM Y-27632 (STEMCELL Technologies #72308) at a 1:1 ratio and added to a 6-well plate. Each well was overlayed with 2 ml MBM medium after Matrigel had solidified in a 37°C and 5% CO_2_ culture incubator for 20 min. ACC organoid culture MBM medium was refreshed once a week. Every 2-4 weeks organoids were passaged by mechanical pipetting Matrigel gently using Dispase in DMEM/F12 media (STEMCELL Technologies #) and several washes with PBS until Matrigel was cleared out. Organoid fragments were then re-suspended in Matrigel and seeded as described above.

### *In vitro* short-term organoid culture cytotoxicity assays

Human ACC patient tumor single cells were embedded in 10 µl of MBM medium with 50% Growth Factor Reduced-Matrigel (Corning #354230) on 384-well white plate at a concentration of 2000 cells per well. After solidification of the Matrigel for 30 min at 37°C, 20 µl fresh MBM medium was added to each well, and the plates were further incubated for 2 days. After the 2 days of pre-culture, cells were treated with 30 µl ADCT-701 and B12-PL1601 for 7 days or with 30 µl SG3199 or other chemotherapeutic drugs for 3 days. For ADCT-701 with or without ABCB1 inhibitors cytotoxicity, NCI-ACC51 and NCI-ACC48 organoid single cells were embedded in Matrigel on 384-well white plates. After 2 days of incubation, cells were treated with different concentration of ADCT-701 or SG3199 combined with or without 1µM valspodar, 10 µM elacridar, and 1µM tariquidar for 7 days or 3 days respectively. For chemosensitivity of NCI-ACC51 and NCI-ACC48 organoids, cells were plated in 384-well white plates as in the previous seeding steps. After 2 days incubation, cells were treated with mitotane, etoposide, doxorubicin, or carboplatin for 3 days. 20 µl of CellTiter-Glo 2.0 reagent (Promega #G9241) was added and the luminescence was quantified with a SpectraMax i3x reader (Molecular Devices).

### IHC staining

4-5 µm sections from formalin-fixed, paraffin-embedded blocks were stained using the Bond Refine polymer staining kit (Leica Biosystems DS9800) for DLK1 antibody (dilution 1:2000, Abcam, ab21682) antibody on the Bond Rx automated staining system (Leica Biosystems) following standard IHC protocols with some modifications. Briefly, the slides were deparaffinized and incubated with E1 (Leica Biosystems) retrieval solution for 20 minutes. Primary antibody was incubated for 1 hr at room temperature and no post-primary step was performed. Cover-slipped slides were scanned with a Aperio CS-O slide scanner (Leica Biosystems). DLK1 immunohistochemistry was scored by a pediatric pathologist. Each case was scored for the most prominent intensity (0-3 with 1 representing equivocal, 2 weak, and 3 strong positive staining) as well as for percentage of staining. A modified H-score was calculated as intensity multiplied by percentage of positively stained cells.

For CD56, Ki67, and synaptophysin immunohistochemistry, auto-stainers Ventana Benchmark Ultra (Ventana, Tucson, AZ), were used. Leica Bond Max (Leica Biosystems, Deerfield, IL) auto-stainer was used for SF1 immunohistochemistry. Validation of these stains was performed on daily clinical laboratory controls by the Anatomic Pathologist on clinical service at the Laboratory of Pathology, National Cancer Institute. A Roche Diagnostics anti-CD56 antibody (rabbit, monoclonal, #760-4596, clone MRQ-42, Indianapolis, IN) was used at a prediluted concentration. An Agilent Technologies anti-Ki67 antibody (mouse, monoclonal, # M7240, clone MIB-1, Santa Clara, CA) was used at a dilution of 1:200. A Perseus Proteomics anti-SF1 antibody (mouse, monoclonal, #PP-N1665-00, clone N1665, Komaba, Japan) was used at a dilution of 1:200. A Roche Diagnostics anti-synaptophysin antibody (rabbit, monoclonal, # 790-4407, clone SP11, Indianapolis, IN) was used at a prediluted concentration.

### Cell lines

Human ACC cell lines CU-ACC1 and CU-ACC2 were obtained from the University of Colorado. CU-ACC cells were cultured in 3:1 (v/v) Ham’s F-12 Nutrient Mixture (Gibco #)–DMEM (Gibco #) containing 10% heat-inactivated FBS (Gemini Bio #100-106), 0.4 μg/mL hydrocortisone (Sigma-Aldrich #H6909), 5 μg/mL insulin (Sigma-Aldrich #), 8.4 ng/mL cholera toxin (Sigma-Aldrich #), 10 ng/mL epidermal growth factor (Invitrogen #), 24 μg/mL adenine (Sigma-Aldrich #) and 1% Penicillin-Streptomycin (Gibco #15140122)]. Human ACC cell line H295R (CRL-2128) was obtained from ATCC and cultured in 1:1 DMEM:F12 (Gibco #11320082) containing 2.5% Nu-Serum (Corning #355100), 1% ITS+ Premix Universal Culture Supplement (Corning #354352), and 1% Penicillin-Streptomycin. Human SCLC cell lines H146, H524, and H1618 were obtained from ATCC. Human SCLC cell line H1436 was obtained from Haobin Chen (Washington University). H146 and H524 cells were cultured in RPMI-1640 (Corning #MT10040CM) supplemented with 10% FBS and 1% Penicillin-Streptomycin. H1618 and H1436 cells were cultured in HITES media DMEM/F12 (1:1) (Gibco #11320082) containing 5% FBS, 1xGlutamaxTM (Gibco #35050061), 10 nM Hydrocortisone, 10 nM beta-estradiol (Sigma-Aldrich #E2758), Insulin-Transferrin-Selenium mix/solution (Invitrogen #41400045), and 1% Penicillin-Streptomycin. All cell lines were cultured at 37°C in a humidified incubator with 5% CO_2_, regularly tested to be mycoplasma-negative (Lonza #LT07-318) and authenticated by STR profiling (Laragen Inc.).

### Flow cytometric analysis

For surface DLK1 expression analysis, human ACC cells, ACC patient tumor cells, ACC PDX cells, and human SCLC cells were harvested and washed with FACS buffer (PBS containing 1% BSA and 0.1% sodium azide). Cells were incubated with the anti-human DLK1 primary antibody (AG-20A-0070-C100 AdipoGen; 1:100 per million cells) at 4°C for 30 minutes in the dark. Cells were washed by FACS buffer. Cells were then incubated with secondary antibody (Invitrogen, #P852; 1:500) at 4°C for 30 minutes in the dark. For surface DLL3 expression analysis, human SCLC cells were collected and washed with FACS buffer. Cells were stained with human DLL3-PE (R&D Systems, #FAB4315P; 10 µl per one million cells) or isotype control antibody (R&D Systems, IC108P; 10 µl per one million cells) at room temperature for 30 minutes in the dark. For surface MDR-1 expression analysis, human ACC cells, ACC patient tumor cells, ACC PDX cells, and human SCLC cells were collected and washed with FACS buffer. Cells were stained with human CD243-PE (Biolegend, #348606; 5 µl per one million cells in 100 µl wash buffer) or isotype control antibody (Biolegend, #400214; 5 µl per one million cells in 100 µl wash buffer) at 4°C for 30 minutes in the dark. Cells were washed and then stained with PI (Biolegend, #421301;1:100) following above antibodies incubation. Living cells were separated as PI negative cells. To semi-quantitate DLK1 or DLL3 cell surface expression in ACC and SCLC cell lines, cell surface molecules of DLK1 or DLL3 per cell were calculated after subtracting background signal from DLK1 secondary antibody alone (Invitrogen, #P852) or DLL3 isotype control antibody (R&D Systems, #IC108P) respectively by BD Quantibrite Beads PE Fluorescence Quantitation Kit (BD Bioscience, #340495) in accordance with the manufacturer’s protocol. Stained cells were acquired on LSR Fortessa (BD Biosciences), and data were analyzed using FlowJo software version 10.8.1. Flow cytometry gating strategies are shown in Supplementary Fig. 11A-C.

### *In vitro* cell line cytotoxicity assays

ACC or SCLC cells were seeded into 384-well white plate at a concentration of 1500 cells per well in 30 µl medium and allowed to adhere overnight. The PCPG cell line, hPheo1, 300 cells per well in 30 µl medium were plated into 384-well white plate for overnight. 30 µl fresh medium containing different concentrations of antibody-drug conjugate (ADCT-701 and B12-PL-1601) or free payload (SG3199) was added to each well and the plates were further incubated for 7 days or 3 days respectively. After 7 days (ADCT-701 and B12-PL1601) or 3 days (SG3199) incubation, 20 µl of CellTiter-Glo 2.0 reagent (Promega #G9241) was added and the luminescence was recorded using a SpectraMax i3x reader (Molecular Devices).

### Imaging flow cytometry

A total of 1 × 10^6^ CU-ACC1 or H295R cells were seeded in each well of six-well plate and allowed to attach overnight. Then, attached cells were incubated with APC-conjugated DLK1 antibody (R&D Systems, a bio-techne brand #FAB1144A) or isotype control antibody (R&D Systems, a bio-techne brand #IC0041A) for 1 hour in 2 ml of cell culture media at 37°C. Then, the cell monolayer was collected and rinsed with cold PBS twice and resuspended. The cellular internalization rate of DLK1 antibody in treated cells was evaluated using an Amnis ImageStreamX Mark II imaging flow cytometry (Luminex, Austin, TX, USA).

### Cell cycle and apoptosis assays

For EdU incorporation studies, cells were processed as per the manufacturer’s instructions (invitrogen #C10634). Briefly, 3 x 10^5^ CU-ACC1 or H295R cells were plated in 6-well plates and allowed to adhere overnight. CU-ACC1 or H295R cells were then treated with 0.02 µg/mL ADCT-701 or B12-PL1601 for 2 or 5 days, respectively. Cells were labeled with 10 µM Click-iT^TM^ EdU in a 37°C and 5% CO_2_ culture incubator for 1 hr. Cells were then fixed and permeabilized. Click-iT™ Plus reaction cocktail was added in cells. Cells were then stained with DAPI for DNA content and detected using a LSR Fortessa cytometer (BD Biosciences) and analyzed using FlowJo software version 10.8.1. For Annexin V staining, 2 x 10^6^ CU-ACC1 or 1.5 x 10^6^ H295R cells were seeded in 6 cm dish and treated with 20 µg/mL ADCT-701 or B12-PL1601 for 1 or 3 days respectively. Apoptosis was detected using an FITC Annexin V Apoptosis Detection Kit with PI (BioLegend #640914) following the manufacturer’s instruction. Briefly, cells were washed with cold PBS containing 1% BSA and 0.1% sodium azide and resuspended in Annexin V binding buffer and stained with Annexin V and PI at room temperature and then analyzed immediately. Annexin V-positive cells were detected using a LSR Fortessa cytometer (BD Biosciences) and analyzed using FlowJo software version 10.8.1.

### Immunoblotting

Cells were lysed in RIPA buffer (Millipore #20-188) supplemented with protease inhibitor (Sigma-Aldrich #11836153001) and phosphatase inhibitors (Sigma-Aldrich #04906837001). Protein concentration was determined by the DC^TM^ Protein Assay Reagents Package Kit (Bio-Rad #5000116). 20 µg of protein lysates were resolved on 4-15% Protein Gel (Bio-Rad #5671084) and transferred to nitrocellulose membrane. The membranes were blocked in 5% blotting grade blocker (Bio-Rad, 1706404XTU) in TBS with 0.1% Tween-20 and then incubated with the indicated primary antibodies. Primary antibodies (1:1000) included DLK1 (CST #2069), phospho-Histone H2A.X (Ser139) (Millipore #05-636), cleaved caspase-3 (Asp175) (CST #9661), cleaved PARP (Asp214) (CST #9541), total NOTCH1 (CST #3608), NOTCH1-ICD (CST #4147), SYP (CST #36406), and CYP17A1 (CST #17334). Primary antibody for detection of α-tubulin (Sigma-Aldrich #T9026) was used at a dilution of 1:1500. Secondary antibodies (1:5000) were from donkey anti-rabbit IgG-HRP (Cytiva #NA934) and sheep anti-mouse IgG-HRP (Cytiva #NA931).

### *In vitro* bystander killing assays

Bystander activity was assessed by co-culturing WT CU-ACC1 and CU-ACC1 DLK1 KO (clone 10) cells at various ratios in white-walled 384-well tissue culture treated plates with complete media. The following day, cells were treated with 1 µg/mL ADCT-701 or B12-PL1601 and incubated in a humidified atmosphere with 5% CO2 at 37°C for 4 days. Cell viability was measured by CellTiter-Glo Luminescent Cell Viability Assay kit (Promega). Bystander activity was also assessed using conditioned media assays in which CU-ACC1 cells were seeded at a density of 1500 cells/well. ADCT-701 was then added in triplicate the next day, in a dose titration ranging from 20 µg/mL to 20 pg/mL. Cells were subsequently incubated for 5 days. On day 5, 30 µL of conditioned media from these plates was removed from each well and transferred to a fresh plate containing CU-ACC1 DLK1 KO (clone 10) or WT CU-ACC1 cells, which were plated 24 hrs previously in 30 µL complete media (final volume 60 µL). These plates were incubated for 5 days before cell viability measurement. Cell viability was determined using the CellTiter-Glo Luminescent Cell Viability Assay kit and data were presented as percent cell viability relative to untreated controls.

### Lentiviral constructs and lentivirus production

For the CRISPR-Cas9 system, a single target sequences for CRISPR interference were designed using the sgRNA designer (https://portals.broadinstitute.org/gppx/crispick/public) and subcloned into the lentiCas9-Blast (Addgene #52962). Viral transduction was performed in the presence of polybrene (5-10 μg/mL, Sigma-Aldrich #TR-1003-G) and cells were centrifuged at 1200 x g for 4 hrs at 30 °C followed by removal of virus and polybrene. After 72 hrs, cells were selected with blasticidin (1-4 μg/mL) for 5 days.

### siRNA-mediated knockdown of DLK1

DLK1-targeting siRNA (siDLK1-1: #4392420 ID s16740, siDLK1-2: #4392420 ID s16738, siDLK1-3: #4392420 ID s16739) and control siRNA (#4390843) were purchased from Invitrogen. CU-ACC1 (5 x 10^5^ cells/well) cells were plated in 6-well plate overnight. Cells were then transfected with siRNA at a final concentration of 30 nM using Lipofectamine RNAiMAX (Invitrogen #13778150). Four days after transfection, whole cell lysates were collected and analyzed by Western blotting.

### Enzyme-linked immunosorbent assay (ELISA)

Cortisol levels were quantified from conditioned media from CU-ACC1 parental and DLK1 KO cells using a cortisol ELISA kit (Enzo Life Science #ADI-900-071). Conditioned media was collected from cells (1 x 10^6^ cells/well) seeded in 12-well plate for 3 days.

### Mice

All animal procedures reported in this study were approved by the NCI Animal Care and Use Committee (ACUC) and in accordance with federal regulatory requirements and standards. All components of the intramural NIH ACU program are accredited by AAALAC International. Seven-week-old female NSG mice were obtained from the CCR Animal Research Program. Seven-week-old female athymic nude mice were purchased from Jackson Labs. All mice were housed in accredited facilities on a 12 hrs light/dark cycle with free access to food and water under pathogen-free conditions.

### *In vivo* efficacy studies

For ACC cell line-derived xenograft models, 2 x 10^6^ CU-ACC1 or H295R cells were subcutaneously injected into the right flank of NSG mice. When the tumor volume reached approximately 100-150 mm^3^, mice were randomized to each group (3-4 mice per group). For ACC PDX models, 164165 and 592788 were obtained from the NCI Patient-Derived Models Repository (PDMR) within the NCI Developmental Therapeutics Program. POBNCI_ACC004 PDX was developed by the NCI Pediatric Oncology Branch. 164165 and 592788 PDX tumor fragments were implanted subcutaneously into the right flank of NSG mice by using a trocar needle. 2 x 10^6^ POBNCI_ACC004 PDX single cells with mouse cell depletion were injected subcutaneously into NSG mice. Recruitment of paired mice in equal numbers to treatment groups was staggered as necessary for any given study. Mice were randomized to each group (5-8 mice per group) once tumors reached 100-200 mm^3^. For SCLC cell line-derived xenograft models, 2 x 10^6^ H524 or H1436 cells were implanted subcutaneously into the right flank of NSG mice. 2 x 10^6^ H146 cells were injected subcutaneously into right flank of athymic nude mice. Tumor-bearing mice were randomized into three treatment groups (4-7 mice per group) once tumor volume reached 100-150 mm^3^. All tumor cells used in vivo were suspended in 100 μL of PBS with 50% Matrigel (BD #356237). Normal saline, B12-PL-1601 (1 mg/kg, diluted in normal saline), and ADCT-701 (1 mg/kg, diluted in normal saline) were intravenously injected into the tail vein on day 0. PDX tumor-bearing mice that initially responded to ADCT-701 treatment were re-treated with ADCT-701 if re-growth tumor volume reached over 100 mm^3^ (otherwise no further doses were administered). Xenograft tumor-bearing mice that did not initially respond to ADCT-701 were re-treated at D7. Body weight and tumor size were measured once or twice weekly respectively, and tumor volume (mm^3^) were calculated with the formula as length x width^2^ x 0.5. Mice were euthanized when tumor volume reached 1500-2000 mm^3^ or 100 days after dosing. Tumors were collected for RNA-sequencing and IHC analysis.

### Quantification and statistical analysis

All statistical tests between groups were unpaired two-tailed Student’s *t*-tests, unless otherwise stated, and *p*-values less than 0.05 were considered statistically significant. Survival analyses were conducted using Cox-proportional hazard models using the R survival package (v3.1.7). Log-rank values were reported for survival analyses. For box plots, the horizontal line represents the median, the lower and upper boundaries correspond to the first and third quartiles and the lines extend up to 1.5 above or below the IQR (where IQR is the interquartile range, or distance between the first and third quartiles).

## Acknowledgments

We also would like to thank the technicians in the CCR Animal Research Program for their support of this study.

## Author contributions

Study conception and design: NYS and NR; Data collection and experiments: NYS; Analysis and interpretation of experiments: NYS and NR; Manuscript writing: NYS and NR. All authors reviewed the results and approved the final version of the manuscript.

## Data and materials availability

Previously published expression datasets re-analyzed in this study can be accessed at GSE10927 and https://portal.gdc.cancer.gov/. Data from Jain et al.^13^ was obtained directly from the authors. Normalized RNA-seq data from the newly generated NCI ACC cohort reported in this study can be found in Supplementary Table 2.

## Notes

### Competing Interest Statement

The authors have declared no competing interest.

