## Supplementary Figures for "Identification of DLK1, a Notch ligand, as an immunotherapeutic target and regulator of tumor cell plasticity and chemoresistance in adrenocortical carcinoma"

#### Supplementary Figure 1

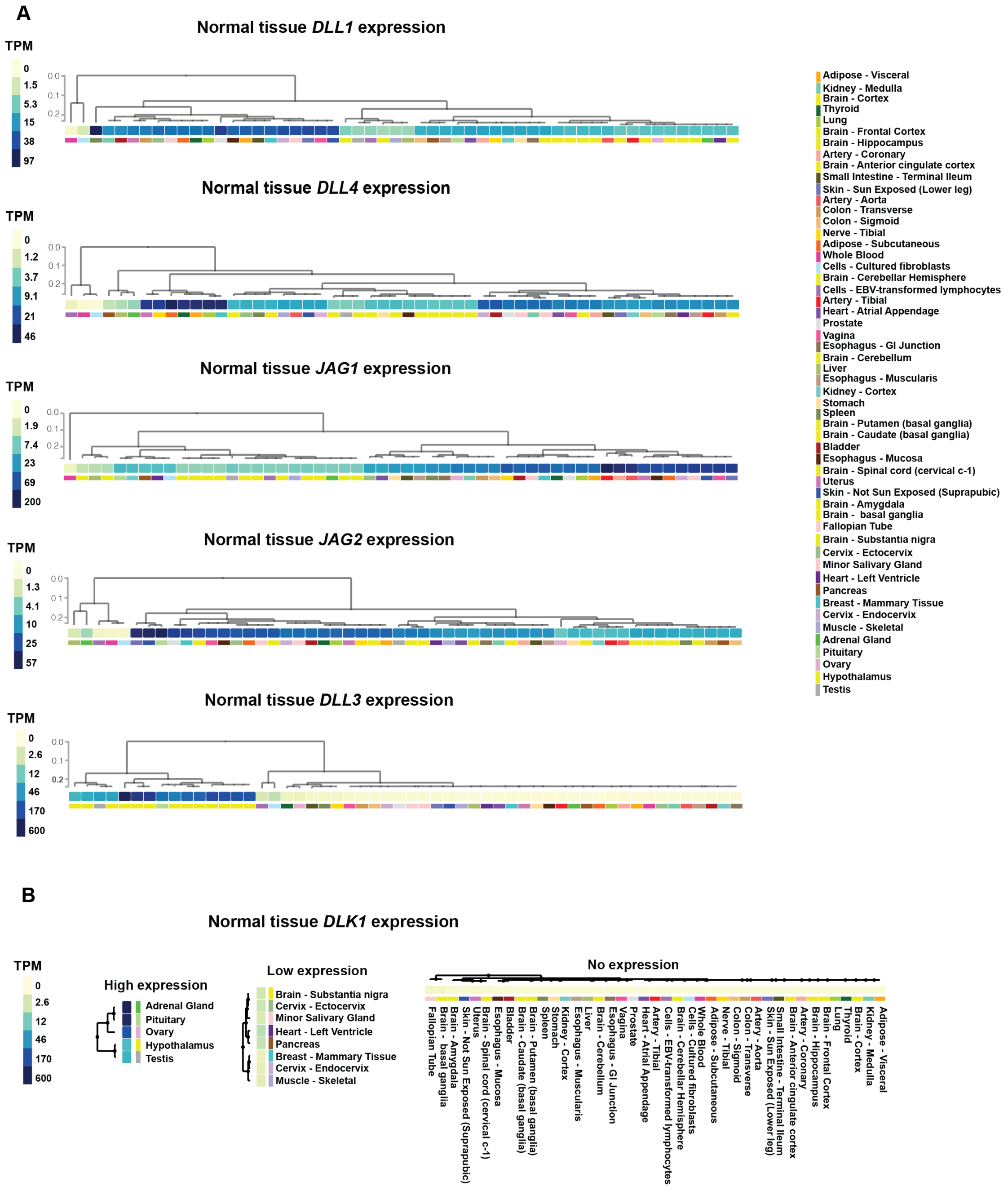


**Supplementary Figure 1. Normal tissue expression of Notch ligands. (A)** Normal tissue expression of *DLL1*, *DLL4*, *JAG1*, *JAG2*, and *DLL3* in the GTEx database. (**B**) Normal tissue expression of *DLK1* in the GTEx database**.**

### Supplementary Figure 2

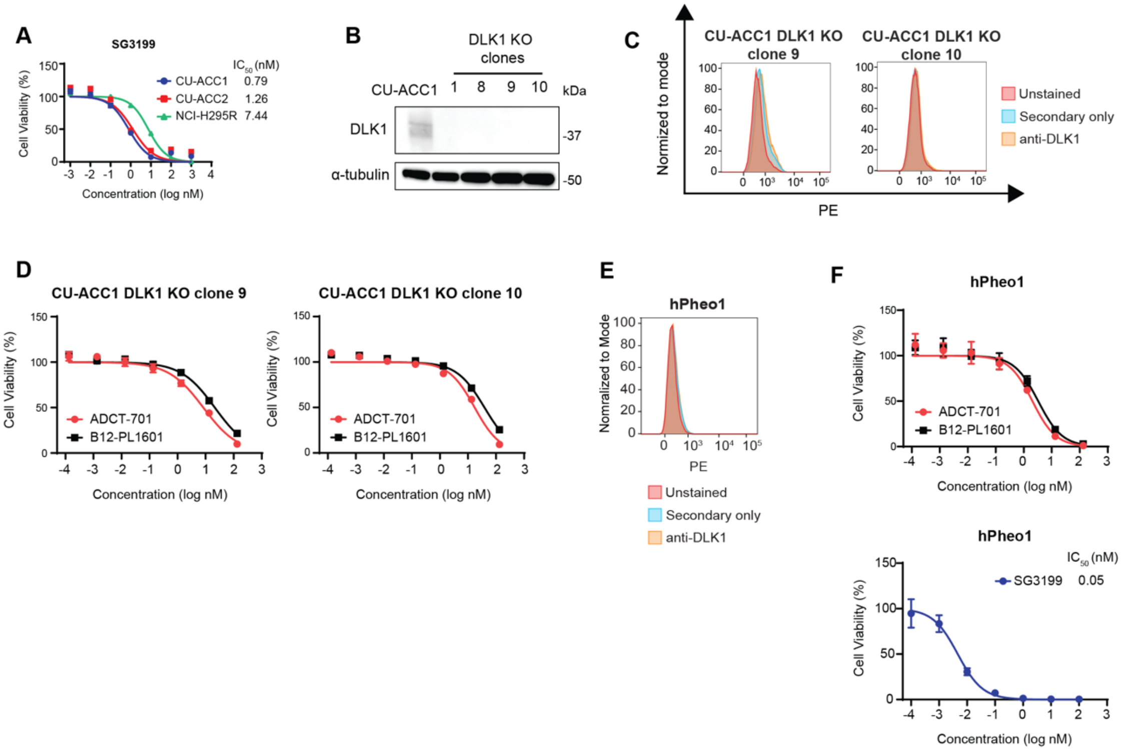


**Supplementary Figure 2. ADCT-701 cytotoxicity is DLK1 dependent. (A)** SG3199 cytotoxicity in CU-ACC1, CU-ACC2, and H295R cells. Cells were treated with SG3199 for 3 days. Each point represents the mean±SEM. **(B)** Immunoblot analysis of DLK1 and loading control (α-tubulin) proteins in CU-ACC1 cells with and without DLK1 KO. Four single-cell KO clones are shown. **(C)** Cell surface expression of DLK1 in two DLK1 KO clones. **(D)** Cytotoxic activity of ADCT-701 in two DLK1 KO clones. **(E)** Cell surface expression of DLK1 by flow cytometry in hPheo1 cell line. **(F)** Cytotoxic activity of ADCT-701 and SG3199 in hPheo1 cell line.

#### Supplementary Figure 3

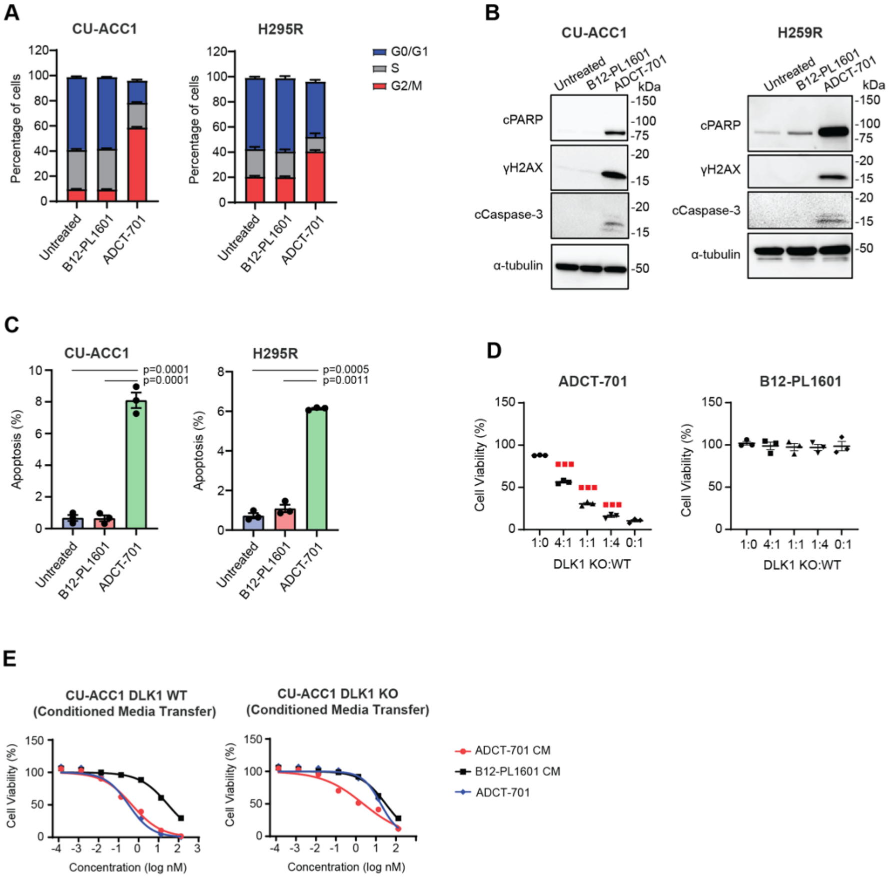


**Supplementary Figure 3. ADCT-701 induces cytotoxicity in DLK1⁺ ACC cell lines through apoptosis and bystander killing.** (**A**) Quantification of the cell cycle in CU-ACC1 and H295R cells treated with 0.02 µg/mL ADCT-701 or B12-PL1601 for 2 or 5 days, respectively. Data are expressed as mean±SEM (n=3 biological replicates). (**B**) Immunoblots showing DNA damage (γH2AX) and apoptosis (cleaved PARP and cleaved caspase-3) in CU-ACC1 and H295R cells treated with 20 µg/mL ADCT-701 or B12-PL1601 for 1 or 3 days, respectively. (**C**) Annexin V apoptosis assay of CU-ACC1 or H295R cells treated with 20 µg/mL ADCT-701 or B12-PL1601 after 1 or 3 days of treatment, respectively. Results represent mean±SEM (n=3 biological replicates). (**D**) Cell viability among different co-culture ratios of CU-ACC1 DLK1 KO:CU-ACC1 treated with 1 µg/mL of ADCT-701 (left) or B12-PL1601 (right) for 4 days. Red lines indicate expected viability with no bystander killing effect. Data are presented as the mean±SEM (n=3 independent experiments). (**E**) CU-ACC1 cells were treated with ADCT-701 or B12-PL1601 for 5 days. Conditioned media was collected and transferred to either CU-ACC1 or CU-ACC1 DLK1 KO cells for 5 days and cell viability was measured. Results represent mean±SEM from 3 independent experiments.

#### Supplementary Figure 4

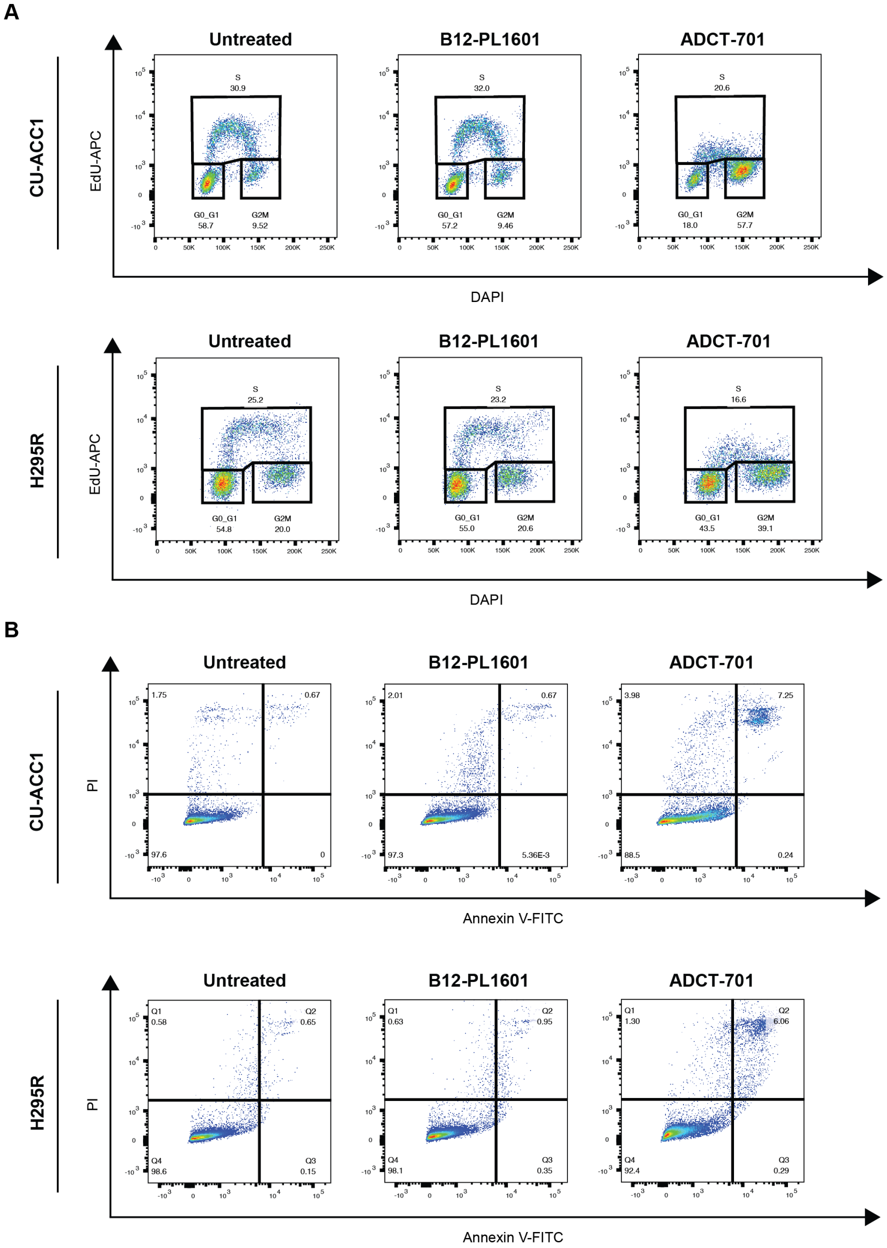


**Supplementary Figure 4. ADCT-701 induces cell cycle arrest and apoptosis among DLK1$^{+}$ ACC cell lines. (A)** Representative flow graphs of cell cycle analysis show cells in S-phase (EdU+) and other phases (G1, and G2M) in CU-ACC1 or H295R cells treated with 0.02 µg/mL ADCT-701 or B12-PL1601 for 2 or 5 days respectively. **(B)** Assessment of apoptosis by flow cytometry of Annexin V and PI double positive CU-ACC1 or H295R cells treated with 20 µg/mL ADCT-701 or B12-PL1601 for 1 or 3 days, respectively.

#### Supplementary Figure 5

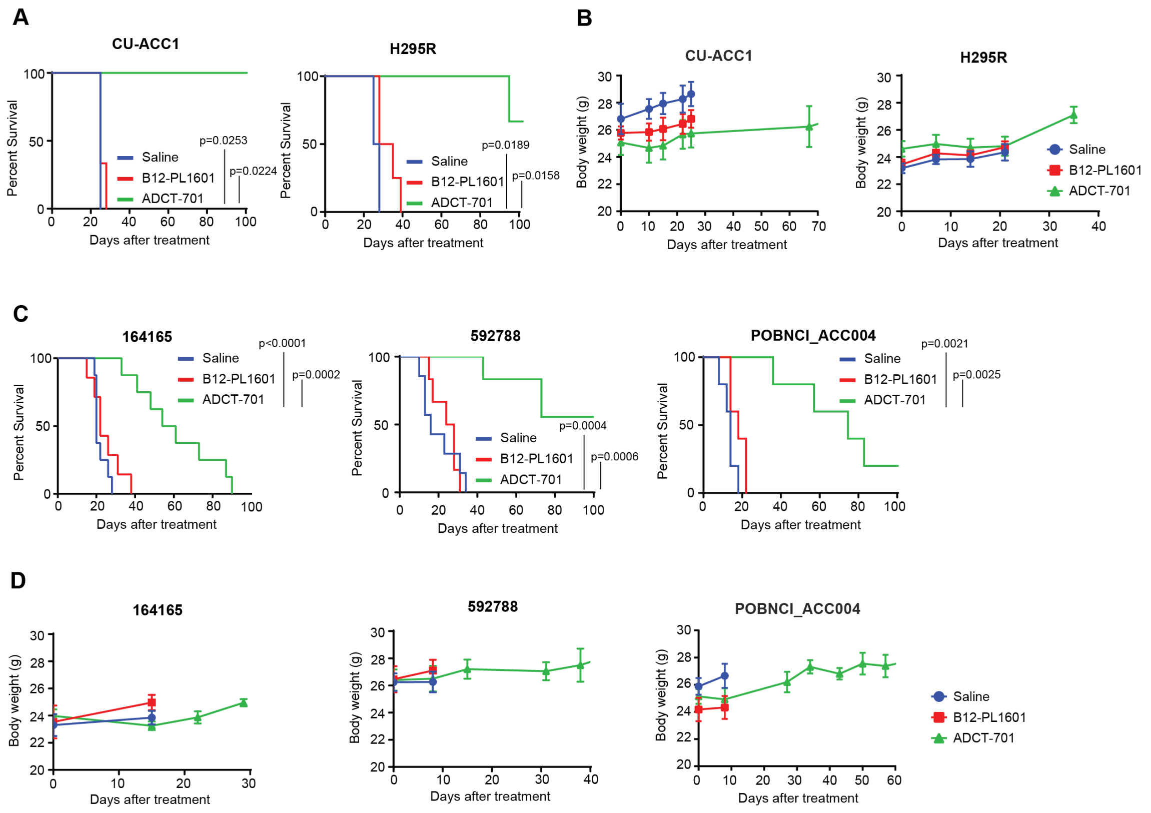


**Supplementary Figure 5. ADCT-701 survival and toxicity in ACC cell-line and patient-derived xenografts. (A)** CU-ACC1 and H295R xenograft survival in mice treated with normal saline, B12-PL1601, or ADCT-701. **(B)** Body weight curves with saline, B12-PL1601, or ADCT-701 treatment in CU-ACC1 and H295R xenografts. (**C**) ACC PDXs 164165, 592788, and POBNCI_ACC004 survival after treatment with saline, B12-PL1601 or ADCT-701. **(D)** Body weight curves with saline, B12-PL1601, or ADCT-701 treatment in 164145, 592788, and POBNCI_ACC004 PDX models.

#### Supplementary Figure 6

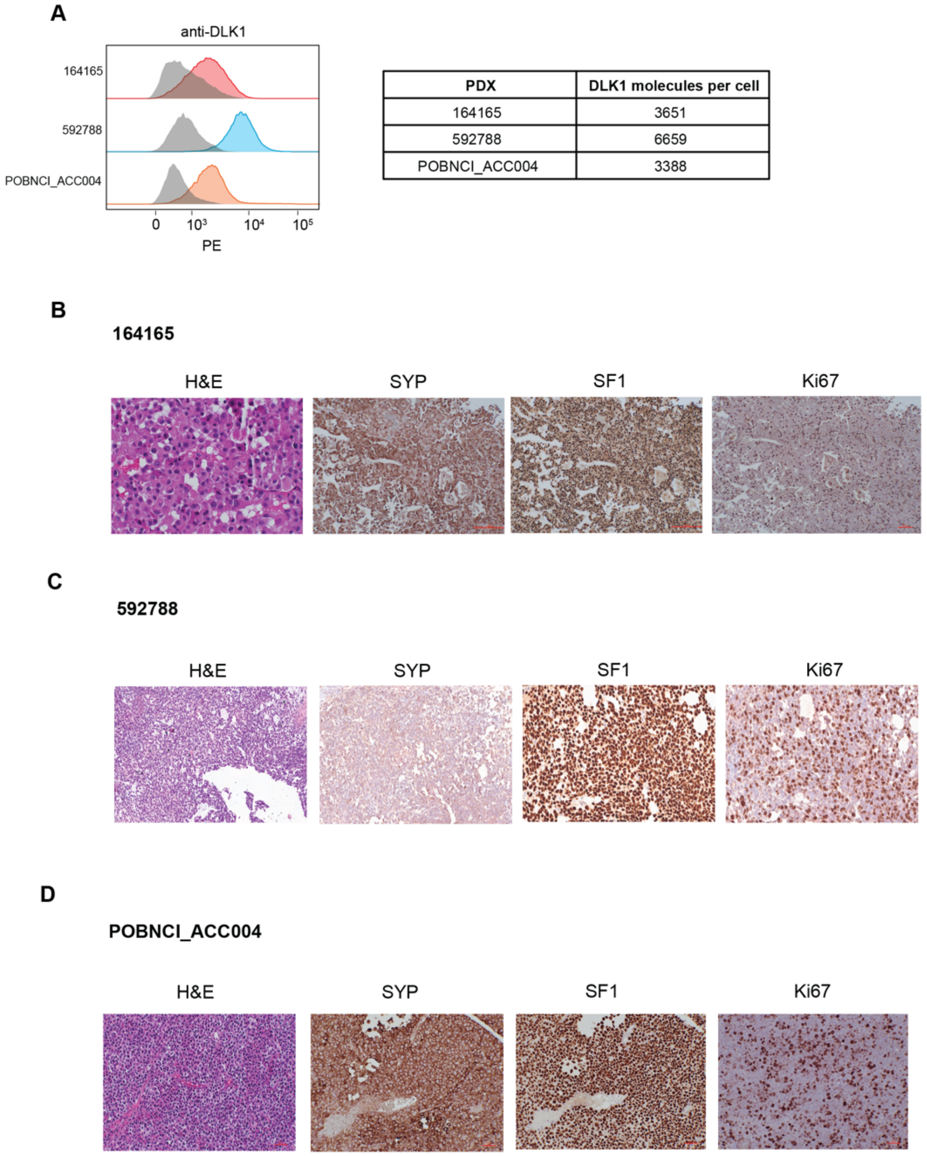


| PDX | DLK1 molecules per cell |
|---|---|
| 164165 | 3651 |
| 592788 | 6659 |
| POBNCI_ACC004 | 3388 |

**Supplementary Figure 6. DLK1 flow cytometry and validation of ACC patient-derived xenograft models. (A)** Cell surface expression of DLK1 and the number of DLK1 molecules per cell among 3 ACC PDX models. H&E staining and immunohistochemistry for NE marker (SYP), adrenal specific marker (SF1) and proliferation marker (Ki67) in **(B)** 164165 PDX, **(C)** 592788 PDX, and **(D)** POBNCI_ACC004 PDX.

#### Supplementary Figure 7

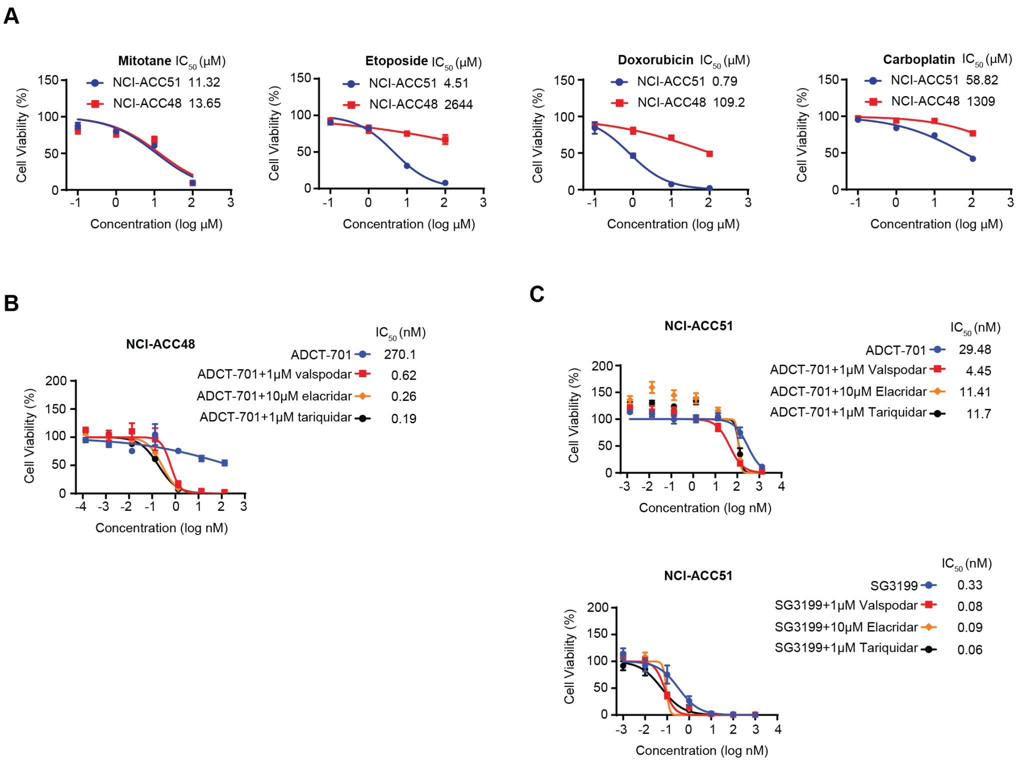


**Supplementary Figure 7. ABCB1 regulates ADCT-701 and chemotherapy resistance among ACC patient-derived organoids. (A)** Cytotoxic activity of mitotane, etoposide, doxorubicin, and carboplatin in two ACC patient-derived organoids (NCI-ACC48 and NCI-ACC51). Cells were treated with chemotherapeutic agents as shown for 3 days. **(B)** ADCT-701 cytotoxicity in the NCI-ACC48 patient-derived organoid with or without ABCB1 inhibitors. **(C)** ADCT-701 and **(C)** SG3199 cytotoxicity in the NCI-ACC51 patient-derived organoid with or without ABCB1 inhibitors. Cells were treated with ADCT-701 or SG3199 combined with or without ABCB1 inhibitors for 7 or 3 days respectively.

#### Supplementary Figure 8

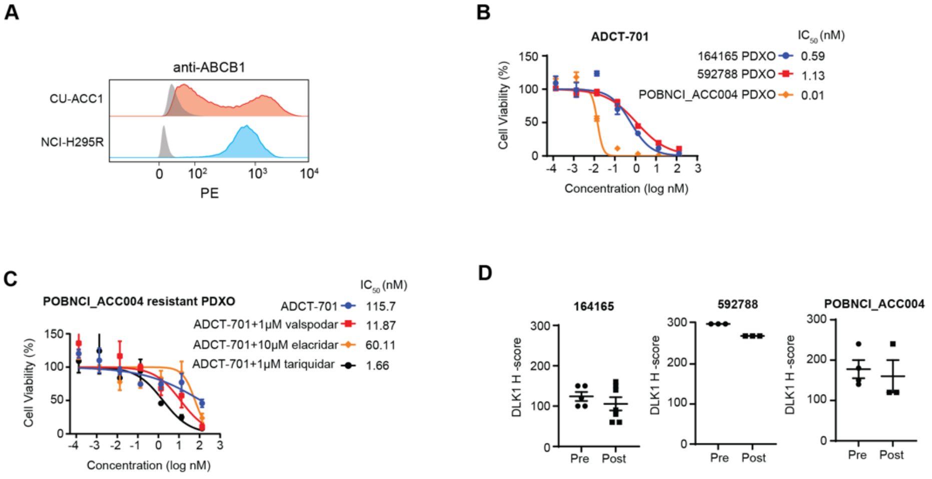


**Supplementary Figure 8. ADCT-701 cytotoxicity in ACC PDX-derived organoids with or without ABCB1 inhibitors. (A)** Flow cytometry histograms of ABCB1 in CU-ACC1 and NCI-H295R cell lines. Shaded gray histograms represent unstained controls for each condition. **(B)** ADCT-701 *in vitro* cytotoxicity in 592788 and POBNCI_ACC004 PDX-derived organoids. (**C**) ADCT-701 cytotoxicity in ADCT-701 resistant PDX-derived organoid treated with or without ABCB1 inhibitors. For panels **B** and **C**, cells were treated with ADCT-701 for 7 days. Each point represents the mean±SEM. (**D**) DLK1 immunohistochemistry in PDX tumors prior to ADCT-701 treatment (Pre) and after resistance to ADCT-701 (Post).

#### Supplementary Figure 9

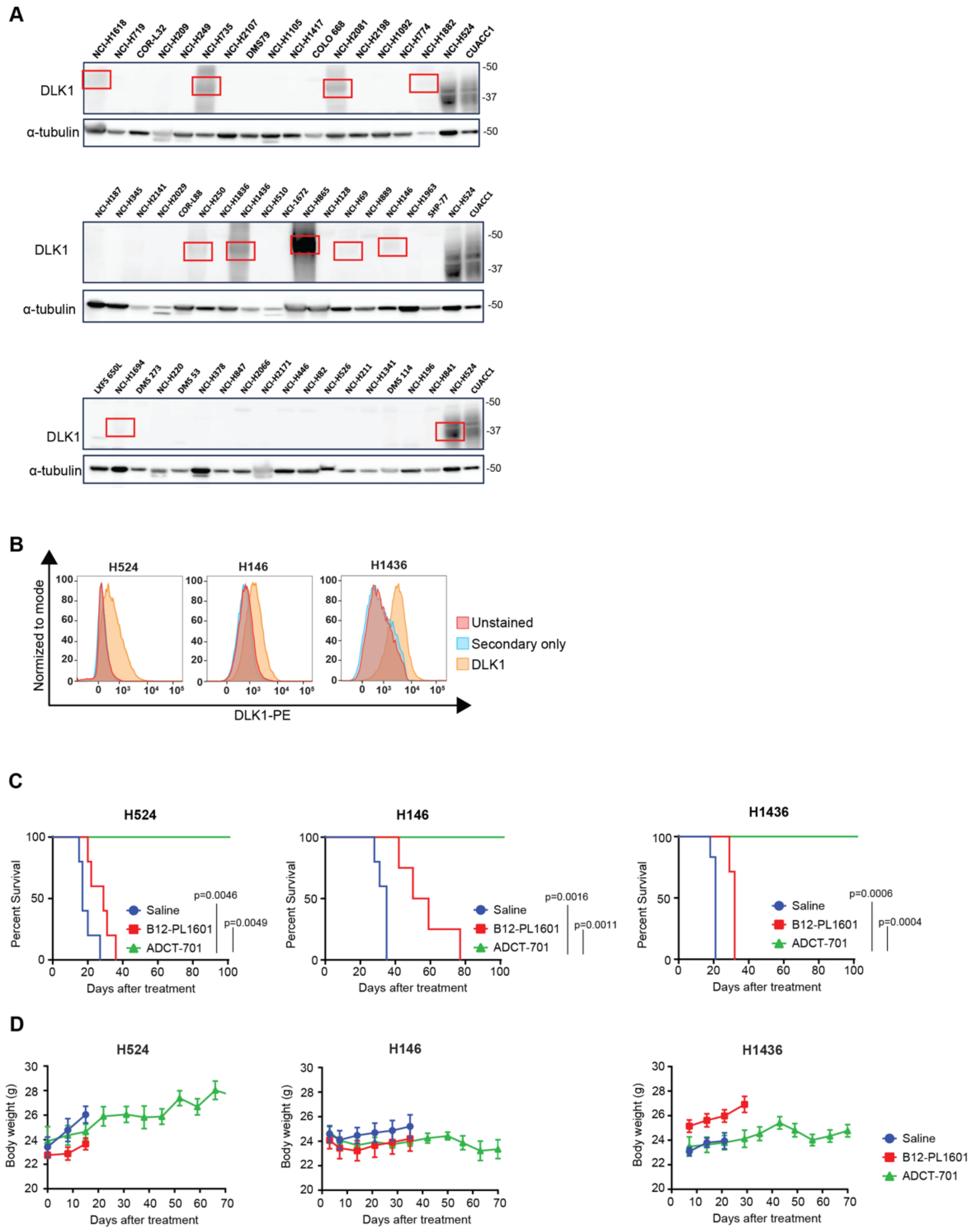


**Supplementary Figure 9. Survival and toxicity of ADCT-701 treatment in DLK1⁺ SCLC xenograft models. (A)** Immunoblot analysis of DLK1 across 51 SCLC cell lines. Cell lines with positive expression of DLK1 are demarcated by the red rectangles. **(B)** Cell surface DLK1 expression by flow cytometry in 3 SCLC cell lines. **(C)** Survival curves of 3 SCLC xenograft models after treatment with saline, B12-PL1601 or ADCT-701 (1 mg/kg) treatment. **(D)** Body weight curves with saline, B12-PL1601, or ADCT-701 treatment in SCLC xenografts H524, H146, and H1436.

### Supplementary Figure 10

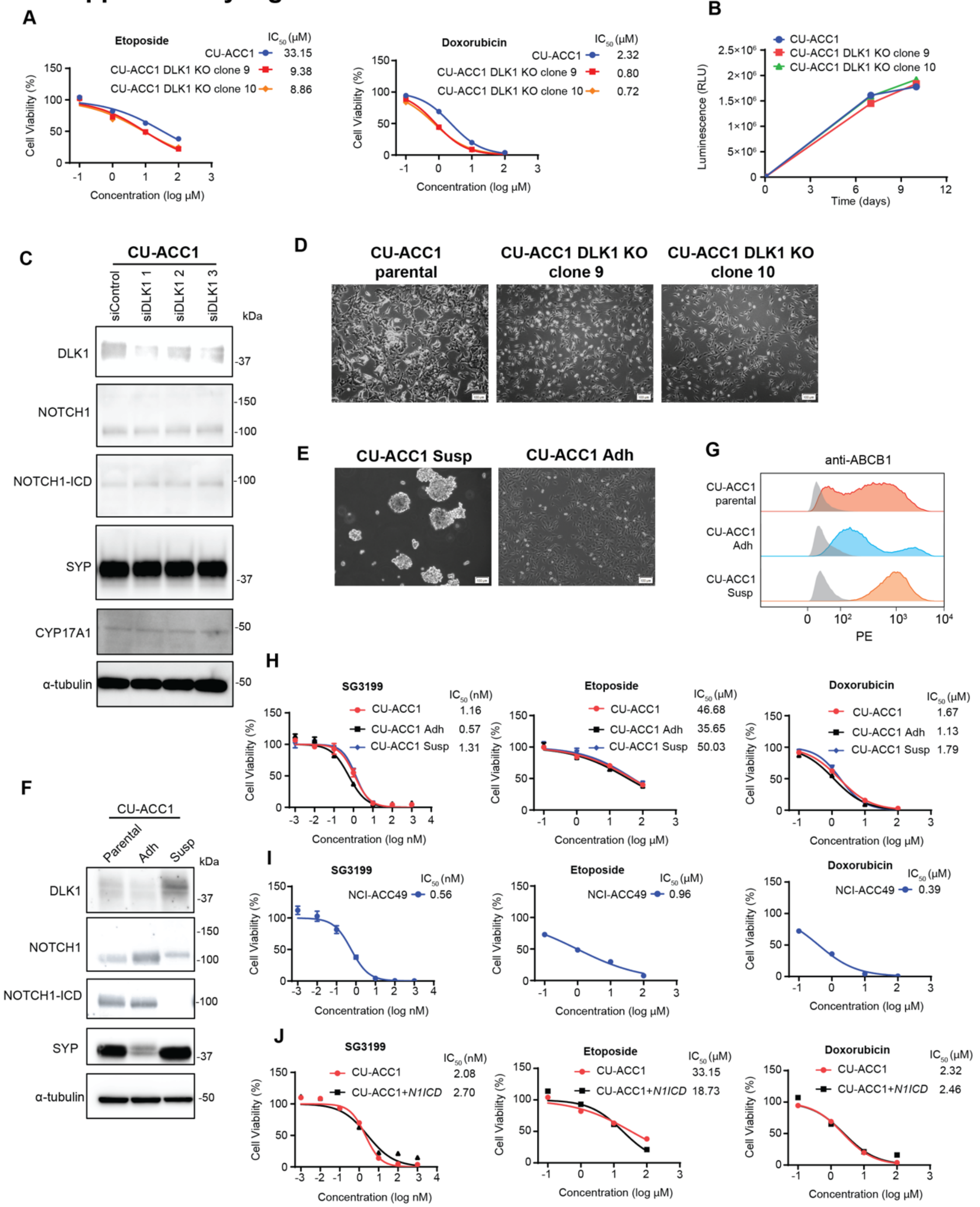

**Supplementary Figure 10. Additional data supporting the relationship between DLK1, NOTCH1 signaling, ABCB1, and chemoresistance in ACC. (A)** Cytotoxic activity of etoposide and doxorubicin in CU-ACC1 parental and DLK KO clones 9 and 10 cells. **(B)** Proliferation of CU-ACC1 parental and DLK KO clones 9 and 10 cells. **(C)** Immunoblot analysis of DLK1, total NOTCH1 and NOTCH1 intracellular domain (ICD), NE marker synaptophysin (SYP), steroidogenic protein CYP17A1, and loading control (α-tubulin) proteins in siControl compared to siDLK1 in CU-ACC1 cells. **(D)** Photomicrographs of CU-ACC1 parental and DLK KO clones 9 and 10 cells. **(E)** Photomicrographs of CU-ACC1 suspension and adherent cells. **(F)** Immunoblot analysis of DLK1, total NOTCH1 and NOTCH1-ICD, SYP, CYP17A1, and α-tubulin proteins in CU-ACC1 parental, adherent, and suspension cells. **(G)** Flow cytometry histograms of ABCB1 expression in CU-ACC1 parental, adherent, and suspension cells. Shaded gray histograms represent unstained controls for each condition. Cytotoxic activity of SG3199, etoposide, and doxorubicin in **(H)** CU-ACC1 parental, adherent, and suspension cells, **(I)** patient-derived organoid NCI-ACC49, and **(J)** CU-ACC1 cells with and without *N1ICD* overexpression.

#### Supplementary Figure 11

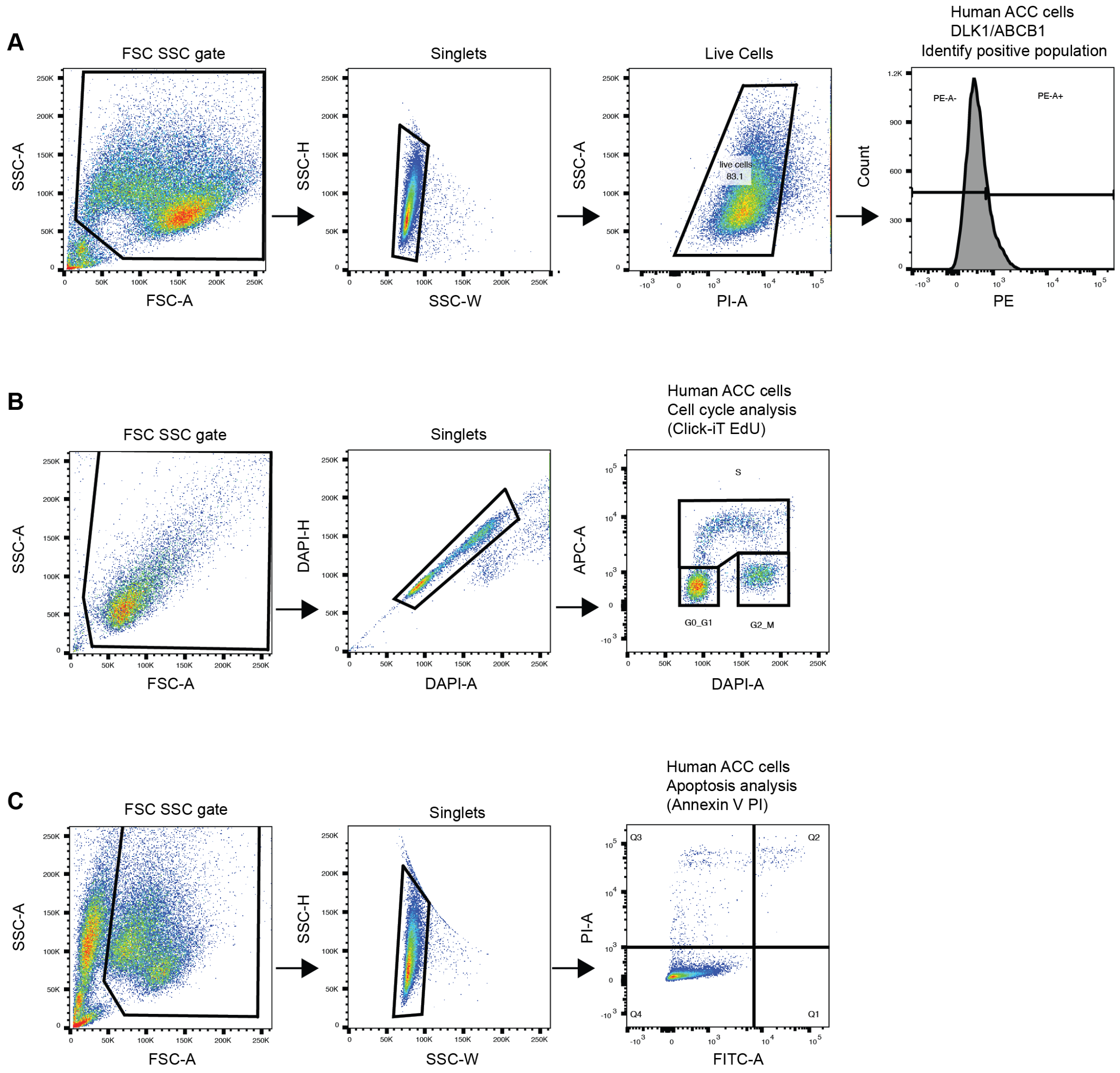


**Supplementary Figure 11. Representative flow cytometric gating strategies. (A)** For cell surface marker analysis, cells were selected based on FSC/SSC profiles and then further gated for singlets and PI-negative populations. **(B)** For cell cycle analysis, cells were analyzed by APC-EdU and DAPI after single cell selection. **(C)** For apoptosis analysis, cells were analyzed by FITC-Annexin V and PI after single cell selection.
